# A dynamic bias in chromatin protein deposition at G-quadruplex sites

**DOI:** 10.1101/2023.09.08.556515

**Authors:** Thamar Jessurun Lobo, Deng Chen, Martijn R. H. Zwinderman, Peter M Lansdorp, Frank J. Dekker, Victor Guryev

## Abstract

Previous studies indicate that genomic loci harboring G-quadruplexes (G4s)—stacked structures that can form in single-stranded DNA—can be linked to epigenetic instability. However, the role of chromatin redistribution and the genome-wide nature of this process need further investigation. Here, we provide experimental evidence that connects G4s to alterations in the deposition of chromatin proteins. We have used metabolic labelling and immunoprecipitation of new and parental proteins in hRPE-1 cells to investigate global chromatin deposition dynamics.

We identify a reciprocal, local bias in chromatin protein deposition at G4 sites favoring the association of parental proteins with the G4 and new proteins with the C4 DNA strand. The deposition bias at G4 sites does not depend on replication directionality and is strengthened by G4 stabilization. Slowing down replication forks upon hydroxyurea treatment reverses the bias, supposedly affected by decoupling between helicase and polymerase. Interestingly, upon combined G4 stabilization and slowing of the replication forks, new proteins exhibit a redistribution pattern similar to G4 stabilization alone, while parental protein redistribution more resembles one after hydroxyurea treatment, hinting at mechanistic differences between parental and new histone distribution. We also report that the genomic distribution of putative quadruplexes is not random and depends on loop size, where G4s with shorter loops have a preference for the DNA strand replicated by leading and G4s with longer loops by lagging strand replication. These findings provide insight into the mechanisms behind G4 occurrence and its role in epigenetic instability and help to improve our understanding of the factors influencing biases in global chromatin protein redeposition.

## Introduction

Single-stranded DNA can form stable stacked structures called G-quadruplexes or G4s, as nearby stretches of guanines bind to each other through Hoogsteen hydrogen bonds (1). Since the discovery of G4s in vitro in 1962, evidence supporting their formation *in vivo* has piled up. The available literature now suggests that G4s affect several cellular processes such as telomere regulation, transcription, RNA splicing, and DNA replication (2, 3). The presence of G4s within most of oncogenic promoters inspired testing them as potential cancer therapeutic targets (4).

Several recent studies have provided indications that G4-formation can also cause epigenetic instability. Thus, deficiencies in DNA helicases FANCJ, BLM, and WRN, involved in DNA repair, were reported to induce a stochastic loss of *BU1A* gene (that has a G4 with the potential to stall the leading replication strand near the gene’s promoter) expression in DT40 chicken cells (5). In the cells deficient in REV1 (a DNA polymerase required for successful replication of G4 DNA) the presence of the G4 correlated with both BU1A expression loss and altered histone modifications at the *BU1A* promoter (6). Interestingly, the study shows that removal of the G4 motif stabilizes *BU1A* expression. Further, when replication stress was induced in these cells by treating them with hydroxyurea (HU) the G4 structure was shown to potentiate expression instability of the BU-1a surface marker and the authors again found alterations in histone modifications at the gene’s promoter (7). Finally, treatment with G4-stabilizer N-methyl mesoporphyrin IX (NMM) decreased BU-1a expression as well, whereas no significant difference was found in cells without the BU-1a G4 (7). These studies suggest that proper replication of G4s is important for the faithful inheritance of the histone code and maintenance of cell-specific expression patterns. They also highlight critical pathways and players that may help to overcome the G4-related perturbations of gene expression. Yet, as few G4-containing genomic loci were investigated so far, it is difficult to evaluate the genome-wide effects of G4 structures on epigenetic stability and the role of chromatin redeposition in this process. To our knowledge, no studies systematically investigated chromatin protein deposition at DNA quadruplex sites. Such investigations would require the application of new experimental techniques that can track histone redistribution throughout the whole genome.

To this end, we recently developed a method that enables direct evaluation of the asymmetry in the distribution of newly synthesized chromatin proteins during DNA replication (8). The method named double-click-seq combines metabolic labeling of new proteins using Azidohomoalanine (AHA, a methionine analog) with metabolic labeling of new DNA using 5-Ethynyl-2’-deoxyuridine (EdU). This allows for subsequent biotinylation and isolation of newly synthesized chromatin proteins and newly synthesized DNA, whereas biotin-(strept)avidin capture allows for stringent washing steps during the isolation process.

Application of this technique on human retinal pigment epithelial (hRPE-1) cells in our previous study demonstrated that newly synthesized chromatin proteins are more often inherited with the lagging DNA strand. Interestingly, this bias is dynamic and can be reverted under stress conditions. Both HU-induced replication stress and the inhibition of DNA damage response pathways resulted in a more frequent association of new chromatin proteins with the leading DNA strand. These observations indicate that the distribution of newly synthesized chromatin proteins over the sister chromatids reflects the difference in the processivity of leading and lagging strand replication.

The effect of DNA replication stress on the redistribution of chromatin proteins prompted us to take a closer look at genomic features that can naturally obstruct DNA synthesis, such as G quadruplexes (G4s). During genome replication, the template DNA is temporarily single-stranded and might thus fold to form G4 structures. Previous studies proposed that G4-associated epigenetic instability results from disturbed chromatin protein deposition as G4 structures in the template DNA interfere with replication (5, 6, 7, 9). Here we aim to find experimental evidence that G4s are indeed connected to alterations in the deposition of chromatin proteins. We applied double-click-seq to hRPE-1 cells to investigate the deposition of newly synthesized chromatin proteins at ∼400,000 genomic regions with experimentally confirmed G4-forming potential (10). We also investigated the effects of G4 stabilization with NMM as well as HU-induced inhibition of replication. In addition, we tracked the distribution of H4 histones with a dimethylation mark at lysine 20 (H4K20me2) as a proxy for the deposition of parental chromatin proteins using SCAR-seq (11). The combination of these two methods (tracking new and old chromatin) provides a detailed genome-wide view of chromatin redeposition dynamics.

## Results

### The double-click-seq method tracks the deposition of labelled histones and structural proteins

One of the limitations of our previous study was uncertainty about the types of proteins that are tracked with the double-click-seq method. To clarify this aspect, we performed a mass-spectrometry assay of proteins that are labelled by AHA and enriched after the EdU-specific click reaction. In the analysis, we checked which proteins are associated with EdU-labelled DNA and also identified peptides containing methionine replaced by L-azidohomoalanine (AHA).

When comparing proteins bound to Edu-clicked DNA to proteins extracted from the same sample that was not subjected to any click reaction, we identify histone H2A as the most enriched by EdU-clicking (P.adj=9.6x10^-17^). The significantly enriched proteins also included histone H2B, actin B and HNRNP proteins (Supplementary Table S1). The peptides with significant evidence for metabolic labeling (Supplementary Table S2)—replacement of methionine by its analog AHA—included histones H4 (3 peptides), H2B (1 peptide, 5 PSMs), actin (5 peptides), and vimentin (7 peptides). These results show that, while histones likely represent the most common chromatin proteins tracked by double-click-seq, part of the signal can also come from other structural nuclear proteins.

### The distributions of parental and newly synthesized proteins at replication initiation zones mirror each other across experimental conditions

In our previous study, we reported that the deposition of new chromatin proteins at replication initiation zones is generally biased towards the lagging strand and that HU-induced replication stress inverts this bias towards the leading strand (8). We now provide evidence for a concerted, complementary effect of replication stress on the deposition of parental and new chromatin proteins. To track the deposition of parental chromatin proteins, we constructed and analyzed SCAR-seq libraries for the same hRPE-1 cell line and experimental conditions (control, hydroxyurea, NMM, and combined treatments, 4 biological replicates per condition, summary of all libraries in Table S3). Using these data, we investigated parental histone deposition (marked by an H4K20me2 modification) at replication initiation zones. We find that parental proteins bind more often to the leading strand in untreated cells, while in cells treated with HU, the distribution is skewed towards the lagging strand (Figure 1). The distribution and effect of G4 stabilization on old chromatin proteins generally mirrors the effect that is seen for new chromatin proteins at replication initiation zones (Supplementary Figure S1). The noteworthy exception from the observed symmetry is a combined treatment with HU and NMM (Figure S1D) where both new and old chromatin proteins show a preference towards the lagging DNA strand, probably due to a more severe blockage of leading strand replication by combined dNTP depletion and G4 stabilization.

**Figure 1.**
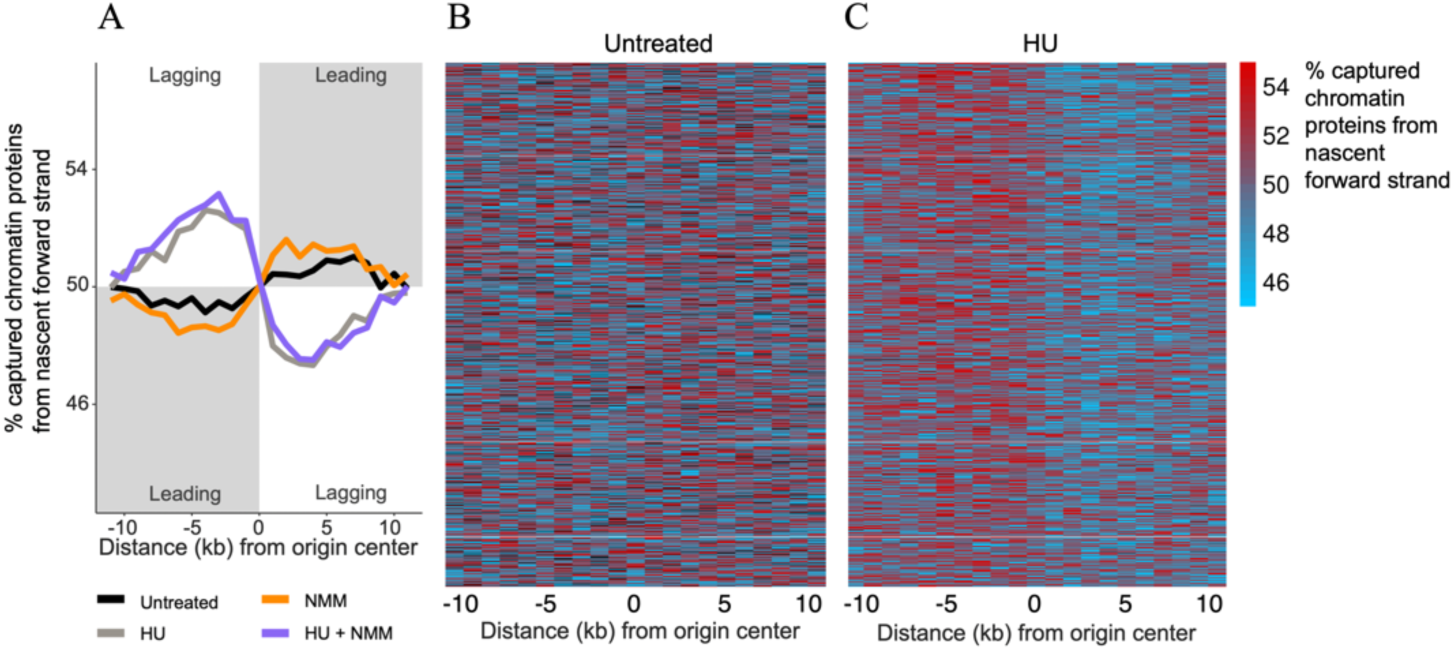
Replication stress inverts the deposition bias of parental chromatin proteins at replication initiation zones. A. The average bias of parental chromatin protein deposition at replication initiation zones under the untreated condition (black line), with HU treatment (grey line), NMM treatment (orange line), and combined treatment (purple line). Separate plots of individual replicates including confidence intervals can be found in Supplementary Figures S2 (double-click-seq, new proteins) and S3 (SCAR-seq, parental proteins). B. Heatmap of parental chromatin protein deposition at replication initiation zones under the untreated condition. C. Heatmap of parental chromatin protein deposition at replication initiation zones in cells treated with hydroxyurea (HU).

### Less new and more parental chromatin proteins bind to template DNA that can form a G4

We used double-click-seq and SCAR-seq to investigate new and parental chromatin protein deposition at 390,129 recently identified sites with experimentally confirmed G4 forming potential (10). A previous study of the *BU-1a* locus (7) only observed epigenetic instability when a G4 was located on the leading strand template. We, therefore, considered the redeposition dynamics of newly synthesized (double-click-seq) and parental (SCAR-seq) proteins at G4 sequences separately for leading and lagging strand replication templates (Figure 2).

**Figure 2.**
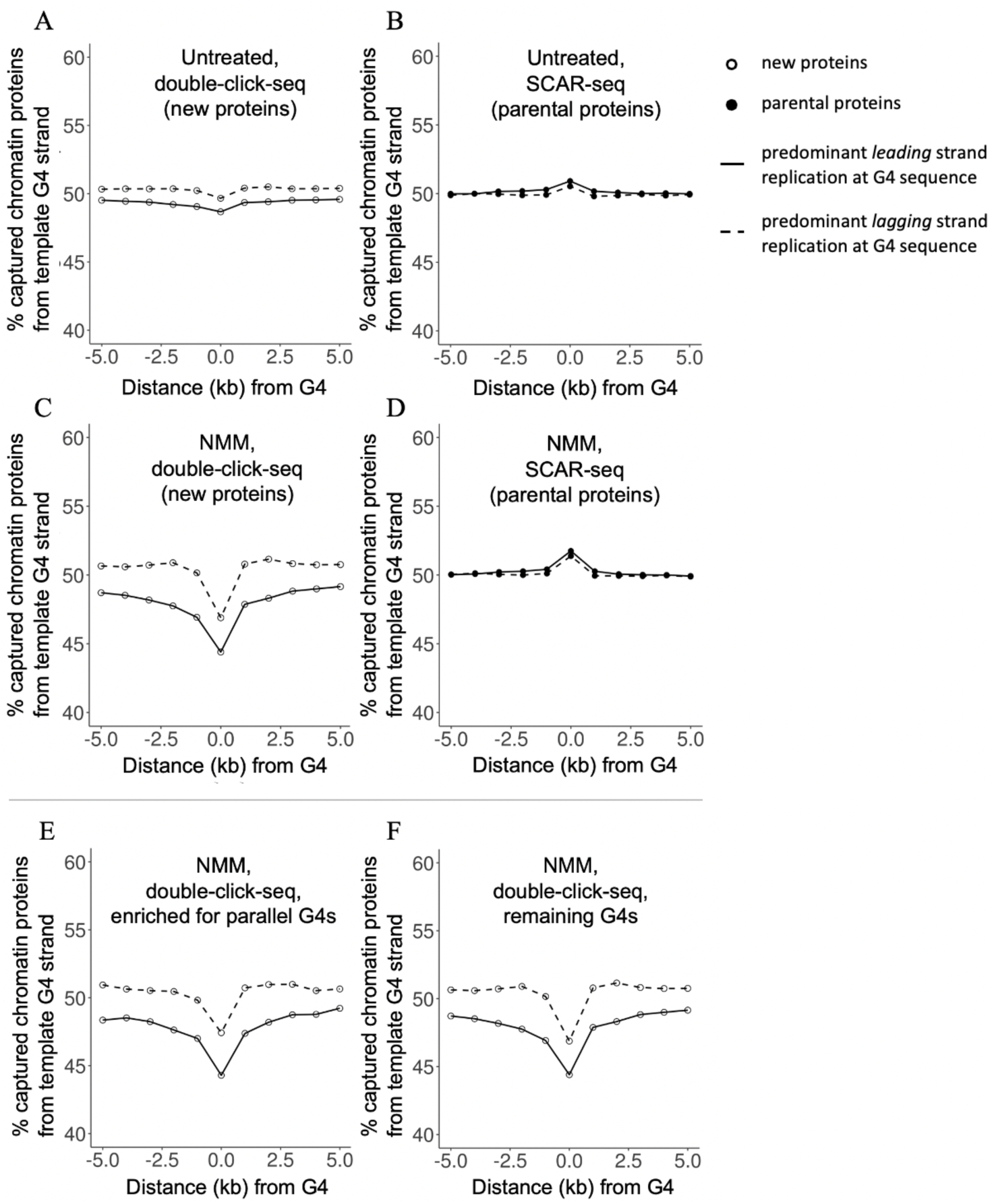
The average deposition of new and parental chromatin proteins at G4 sequences. We determined the average deposition separately for G4 sequences that could interfere with predominant leading (continuous line) and lagging strand replication (dashed line). The deposition was calculated as the percentage of captured chromatin proteins bound to the G4 template strand. A & B. The average deposition of new and parental chromatin proteins in untreated cells. C & D. The average deposition of new and parental chromatin proteins in cells treated with NMM. E. The average deposition of new chromatin proteins at parallel G4 sequences in cells treated with NMM. F. The average deposition of new chromatin proteins at remaining G4 sequences in cells treated with NMM. Separate plots of individual replicates including confidence intervals can be found in Supplementary Figure S4 (double-click-seq, new proteins) and S5 (SCAR-seq, parental proteins).

We identified a reciprocal bias in the deposition of chromatin proteins at G4 sequences, with newly synthesized proteins binding more often to the C4 strand and parental chromatin proteins binding more often to the G4 strand (Figures 2A, 2B and Supplementary Figures S7A and S8A). Interestingly, the effect is observed regardless of the predominant type of replication (leading or lagging strand) at the G-quadruplex sites.

### A G4 stabilizing agent strengthens the redistribution bias

Next, we aimed to confirm that the observed bias in the deposition of chromatin proteins at G4 sequences is a consequence of G4 structure formation. To that end, we investigated the effect of G4 stabilization under N-methyl mesoporphyrin IX (NMM, Figures 2C, 2D and Supplementary Figures S7B and S8B) treatment. Similarly to the untreated cells, we observe less frequent binding of new and more frequent binding of parental chromatin proteins at G4 sites of the template DNA, irrespective of the replication strand.

Interestingly, treatment with the G4 stabilizing agent substantially enhances the redistribution bias, which becomes over four times stronger for new (compare Figure 2C versus 2A) and roughly twice as strong for parental proteins (compare Figure 2D versus 2B). The fact that NMM treatment increases the earlier observed effect at G4 sequences indeed points to G4 structure formation as a probable cause for the biased protein redistribution effect.

### The effect of NMM treatment is not limited to parallel G4 sequences

Intramolecular G4s can be categorized into three groups based on their conformations: parallel when all strands point in the same direction, antiparallel when two strands have opposite orientations than the other two, and hybrid when one strand has a different direction than the other three (12). Previous studies found that *in vitro* NMM specifically stabilizes G4s in the parallel conformation (12, 13). Furthermore, NMM has been reported to be capable of induction of a conformational change from hybrid G4 to parallel G4 (12). But support for the stabilization of antiparallel G4s by NMM is lacking.

We evaluated if the bias-strengthening effect of NMM treatment was limited to G4s known to fold in a parallel conformation. To this end, we divided the G4 sequences into two groups based on their potential to form parallel G4 structures and plotted the average deposition rate of new and parental chromatin proteins in each group (Figure 2E, F and Supplementary Figure S6). When we compare the two groups, we do not observe a stronger effect at G4 sequences that predominantly fold in a ‘parallel’ conformation. This indicates that NMM treatment leads to a greater deposition bias at a broad range of G4 structures.

### Replication stress caused by hydroxyurea treatment reverses chromatin redeposition bias at G4s

Another common way to interfere with replication and induce epigenetic instability is hydroxyurea treatment (7). The proposed mechanism involves depletion of the nucleotide pool through inhibition of ribonucleotide reductase (14), leading to stalling of DNA replication forks. That, in turn, is thought to cause decoupling between the replicative helicase and polymerase, which is associated with runs of single-stranded DNA (15, 16). As a result, hydroxyurea treatment-induced ssDNA formation can be an alternative way to induce G4 formation and affect the epigenetic stability of dividing cells.

In our previous study (8) we showed that treatment with hydroxyurea reverses the deposition dynamics so that new chromatin proteins prefer to associate with the leading rather than lagging strand.

When focusing on G4 sites, hydroxyurea treatment results in a slight preference for new proteins to associate with the G4 strand irrespective of replication directionality (Figure 3A, Supplementary Figure S7C). The matching trend is indistinguishable for parental proteins (Figure 3B, Supplementary Figure S8C), where deposition at G4 sites does not seem to differ from the neighboring regions. This may be due to the generally lower magnitude of changes scored by SCAR-seq, compared to the double-click-seq method. Alternatively, HU-induced slowing of replication forks might act differently on the distribution of parental and new proteins. Because HU and NMM were previously found to cause expression loss of BU-1a in a synergistic manner (7), we have also tested their combined effect on the deposition of chromatin proteins at G4 sequences.

**Figure 3.**
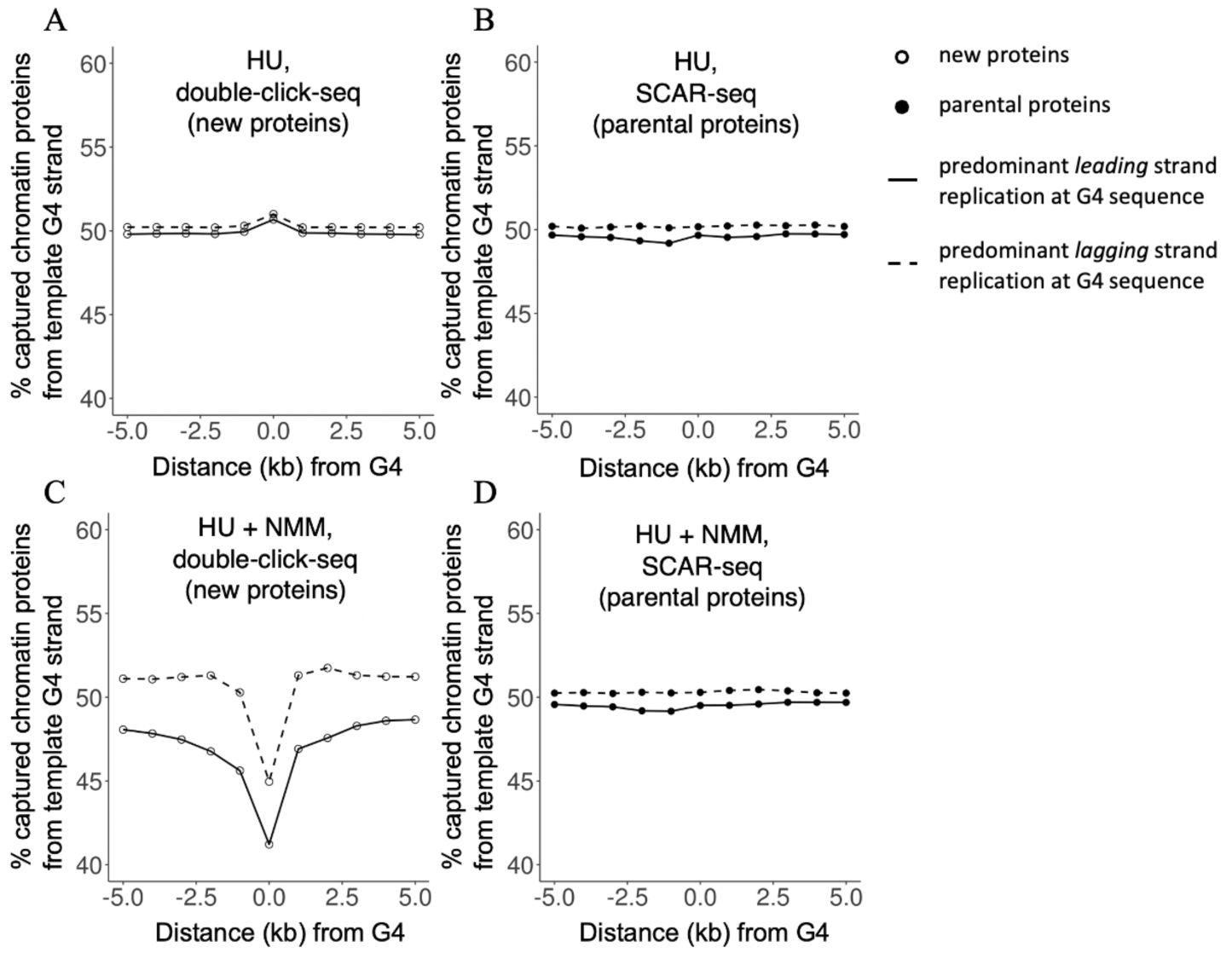
Hydroxyurea treatment affects new and parental chromatin protein deposition in distinctive ways. We determined the average deposition of chromatin proteins separately for G4 sequences that could interfere with predominant leading (continuous line) and lagging strand replication (dashed line). The deposition was calculated as the percentage of captured chromatin proteins bound to the G4 template strand. A & B. The average deposition of new and parental chromatin proteins in cells treated with HU. C & D. The average deposition of new and parental chromatin proteins in cells treated with HU and NMM. Separate plots of individual replicates including confidence intervals can be found in Supplementary Figure S4 (double-click-seq, new proteins) and S5 (SCAR-seq, parental proteins).

Interestingly, we see that the addition of HU further strengthens the effect on the distribution of new chromatin proteins observed in cells treated with just NMM (Figure 3C and Supplementary Figure S7D). Thus, the effect of HU on the deposition of new chromatin proteins differs depending on whether G4s are stabilized or not. When the cells are solely treated with HU, the redeposition bias observed for G4s in untreated or NMM-treated cells inverts. When cells are treated with hydroxyurea and NMM, the bias in the redistribution of new chromatin proteins characteristic of NMM treatment increases.

On the other hand, we do not observe a differential effect of combined treatment with HU and NMM on the deposition of parental chromatin proteins (compare Figure 3D versus Figure 3B and Supplementary Figure S8D versus S8C). This might suggest that combined fork stalling and G4 stabilization disrupts the equilibrium between the deposition of newly synthesized and parental chromatin proteins.

### Fine-mapping of the chromatin protein distribution bias at G4s reveals the local nature of the effect

The effect on the redeposition of chromatin proteins is strongest at the G4 itself and fades out quickly within several kilobases from it (Figures 2,3). To assess the symmetry and local magnitude of the effect we evaluated the biases at sub-kilobase level (Figure 4). Despite much more variation in the data due to lower data density at a higher resolution, we could make a couple of interesting observations. First of all, the bias is most pronounced locally, within 200-300 base pairs from the quadruplex sequence. Secondly, the summit of the distribution is likely shifted upstream of the G4 for both lagging and leading DNA strands. Interestingly, the fragment density distribution of double-click-seq and SCAR-seq libraries shows a characteristic increase of fragments bound by labeled proteins positioned slightly upstream of G4s (Figure 5). The density profiles are complemented by a lower fragment density next to the G4s, and a fading oscillation (double-click-seq, Figure 5A) with a period that is comparable to that observed in nucleosome phasing.

**Figure 4.**
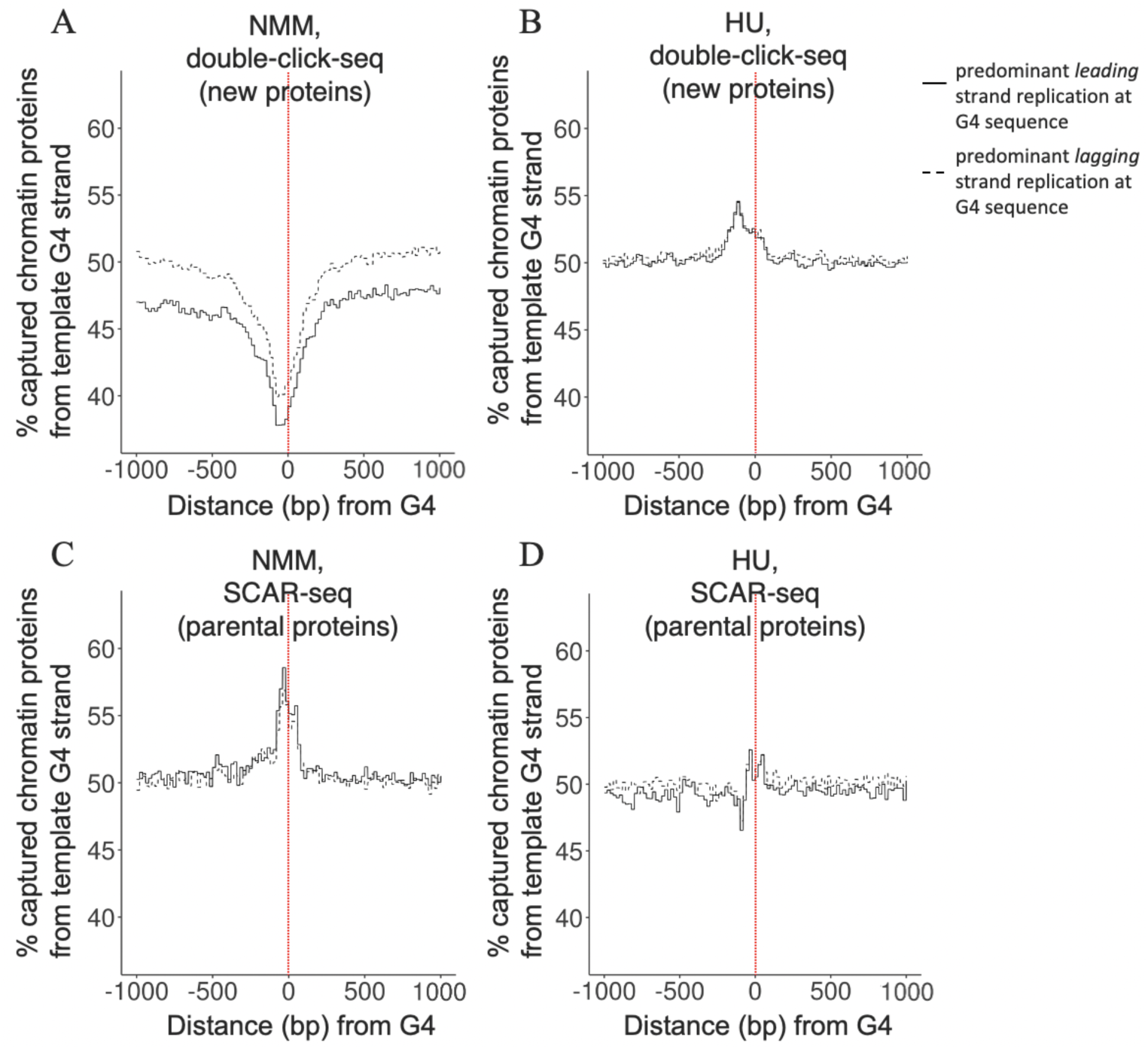
The bias in the deposition of chromatin proteins is strongest immediately at the G4 site. We determined the average deposition separately for G4 sequences that could interfere with predominant leading (continuous line) and lagging strand replication (dashed line). The deposition was calculated as the percentage of captured chromatin proteins bound to the G4 template strand. The vertical red dotted line at x=0 indicates the center of the G4 sequence. A. The average deposition of new chromatin proteins in cells treated with NMM. B. The average deposition of new chromatin proteins in cells treated with HU. C. The average deposition of parental chromatin proteins in cells treated with NMM. D. The average deposition of parental chromatin proteins in cells treated with HU.

**Figure 5.**
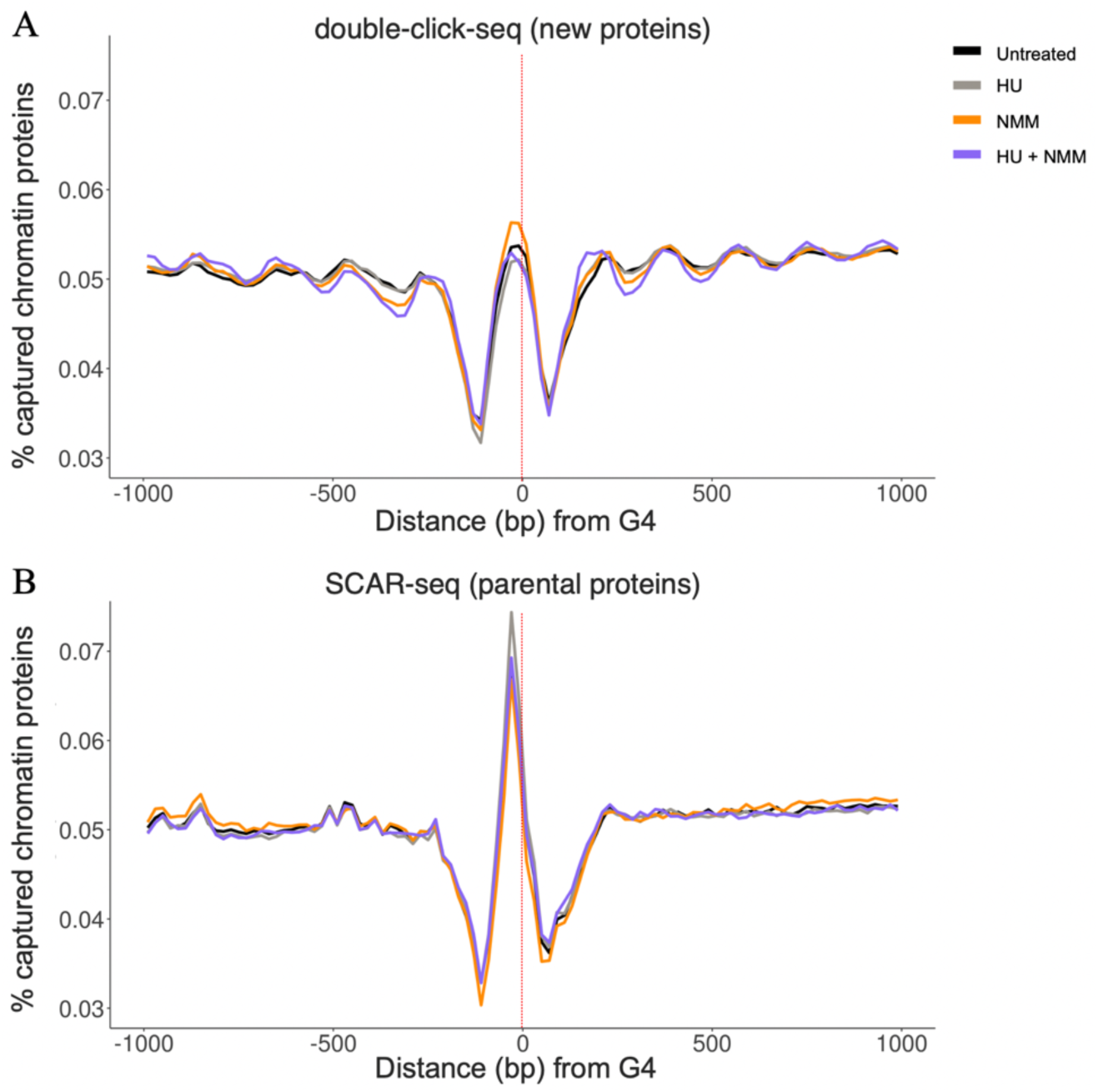
The density of DNA fragments bound to labeled new and parental chromatin proteins relative to G4 regions. The signal was summed across replicates, converted to percentages, and averaged per 20bp windows. A. The average occupancy of new proteins. B. The average occupancy of parental proteins

These results show that the effect of G4s on chromatin protein redistribution is most pronounced locally, within several hundreds of base pairs, slightly upstream of G4, with a similar dynamic for leading and lagging strands. The phasing of DNA fragments at G4s might indicate that G4s are preferentially bound by parental histones under control or G4 stabilizing conditions.

### The distribution of putative quadruplexes and G4 motifs is biased towards genomic regions replicated by lagging and leading strand replication

As we discover new roles for DNA quadruplexes in gene regulation, genome, and epigenome stability, we also note that they might more frequently occur in specific genomic regions. Indeed, previous studies have shown the enrichment of putative G4s in promoters and splice junctions (10, 17). We asked whether such a bias is present concerning the preferred DNA template strand that is used to replicate the G4. The occurrence of G4 sequences identified by the G4-seq method is not random with respect to their mode of replication (leading or lagging strand). The putative G4 sequences are significantly more often positioned to interfere with lagging strand replication (n=170,170 versus n=158,596). This preference for G4 sequences to reside at the lagging strand is relatively mild (7.3% more G4s on the lagging than on the leading strand), but might potentially reflect an adaptation for faster genome replication as G4s are thought to only stall the replicative helicase at the leading strand (18).

Interestingly, the distribution of computationally recognized G4 motifs among DNA strands replicated through leading and lagging strand synthesis depends on the size of the loop between G-stretches that is allowed by the search algorithm. G4 motifs with a shorter loop size (1-6 nucleotides) are biased toward the leading strand while setting this parameter to 7 and more nucleotides increases the proportion of G4s located on the lagging strand (Figure 6). This observation hints that the distribution of G4s at the leading and lagging DNA strands is non-random and potentially depends on G4 conformation and/or stability. A follow-up study that would investigate the relation between G4 type, their position inside genes, and replication direction should unravel the forces behind these distribution biases.

**Figure 6.**
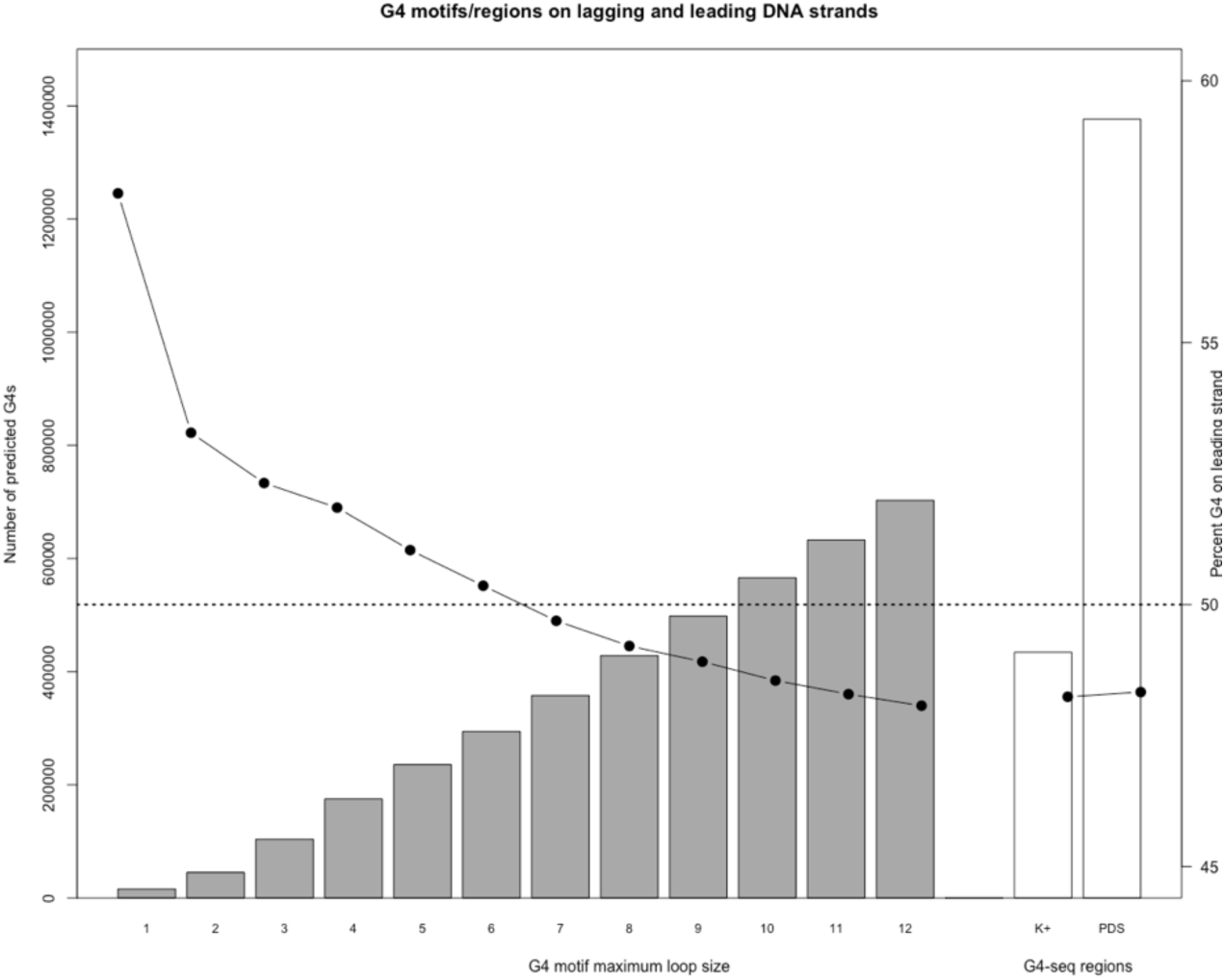
Preference for the leading or lagging strand depends on G4 motif/region definition. The bar plot shows the number of G4s (motifs, gray bars) predicted or G4 regions scored (G4-Seq, where quadruplex structures are stabilized by addition of K^+^ ions or PDS, white bars) in the human genome. The line plot shows the percentage of G4s predicted to interfere with leading strand replication (right axis). The expected leading strand ratio (50%) is indicated by the dotted line.

## Discussion

In this study, we aimed to investigate new and parental chromatin protein deposition at G4 sequences. To do so, we employed two complementary methods to estimate the deposition bias of chromatin proteins in DNA replication. Newly synthesized chromatin proteins were tracked by a metabolic labeling technique (double-click-seq) while, parental chromatin proteins were tracked by the ChIP-based SCAR-seq method. These two methods generally exhibit an opposite bias for paternal and new proteins, compatible with the expectation of complete redistribution (e.g. if one strand receives more new proteins, it receives fewer parental ones). This combination of methods allows us to get a more comprehensive picture of parental and new chromatin proteins distribution upon native and perturbed DNA replication.

Our study represents the first direct comparison of methods for deposition bias identification that rely on metabolic and epigenetic labeling of chromatin proteins. The results show that metabolic labeling (double-click-seq) can identify a higher magnitude of deposition changes at G4 sites. This might be a result of different enrichment efficiency as biotin-streptavidin binding used in double-click-seq is several orders of magnitude stronger than binding of antibody to its target when using SCAR-seq method. Next to that, epigenetic modifications are more dynamic and can be re-written. Finally, the difference can result from variation in chromatin protein composition as our SCAR-seq data tracks the distribution of parental H4 histones, while the double-click-seq method can track a broad spectrum of chromatin proteins with methionine residues.

To experimentally evaluate the protein composition tracked by double-click-seq, we performed a mass-spectrometry analysis of proteins that associate with DNA after an EdU-specific click. The proteomic results showed that, expectedly, histone proteins head the list of proteins that associate with replicated DNA. Next to that, a protein modification search provided evidence for the presence of AHA replacing methionine in the labeled histones. Such a major contribution of histones to the double-click-seq method is also supported by a characteristic phasing pattern observed around G4s (Figure 5), although we do not observe a similar phasing when centering on transcription start sites (Supplementary Figure S9). It is likely that a sizeable fraction of chromatin proteins assessed by the double-click-seq technique represents structural nuclear proteins such as actin.

The combination of SCAR-seq and double-click-seq methods allowed us to monitor chromatin protein redeposition around DNA quadruplex sites. The current literature suggests that G4-formation can likely affect DNA replication and cause epigenetic instability. The latter supposedly is a consequence of the former, as blocked DNA replication at unresolved G4s on the template strand can disturb chromatin protein redeposition at the replication fork. However, we are not aware of any studies investigating chromatin protein deposition at G4 sites. Therefore, we used double-click-seq and SCAR-seq to examine the deposition of new and parental chromatin proteins respectively. We employed human hRPE-1 cells to quantify the redeposition at ∼400,000 sequences with experimentally confirmed G4-forming potential.

We have identified a reciprocal bias in the deposition of new and parental chromatin proteins at these G4 sequences. The newly synthesized proteins bind more often to the C4 strand and parental ones more often associate with the G4 strand. We notice that the distribution bias is similar, regardless of the replication directionality (whether G4 forms at the leading or lagging strand template). This observation opposes the idea that lagging strand replication is not affected by hurdles, such as G4s, due to its ability to reprime (19). The observation of a consistent bias at G4 sites also argues against the possibility that coating of the lagging strand template with RPA can efficiently prevent G4 formation (20). Our conclusions are in line with the results of a previous study in yeast which reported that G4 helicase PIF1 is essential for fast lagging strand replication of template G4 sequences (21).

G4-stabilization with NMM strengthens the observed effect on chromatin protein deposition at G4 sequences. This indicates that G4 structure formation indeed underlies the biased redeposition. Surprisingly, we find that the average bias at a set of G4 sequences enriched for estimated parallel G4 structures is comparable to the bias at the remaining G4 sequences. Currently available evidence from *in vitro* studies only supports the stabilization of parallel G4 structures by NMM (12, 13). Our data, however, suggest that either NMM might also sustain other G4 topologies, be able to convert non-parallel structures to parallel, or that classification of G4 structures from their nucleotide motifs (22) needs further improvement.

During replication, the replicative helicase composed of Cdc45, MCM2-7, and GINS (CMG complex) is thought to move along the leading strand template (23). Accordingly, a previous work using Xenopus egg extracts (18) has shown that the CMG only stalls at G4 sequences on the leading strand template. However, this does not mean that G4 structures on the lagging strand template are not detected by the CMG as it passes by. Recent evidence supports the recognition of G4 structures by a CMG associated protein Timeless (24). We propose that, in both untreated cells and cells treated with NMM, G4 formation on either strand during DNA replication might immediately be recognized and acted upon by the CMG (Figure 7).

**Figure 7.**
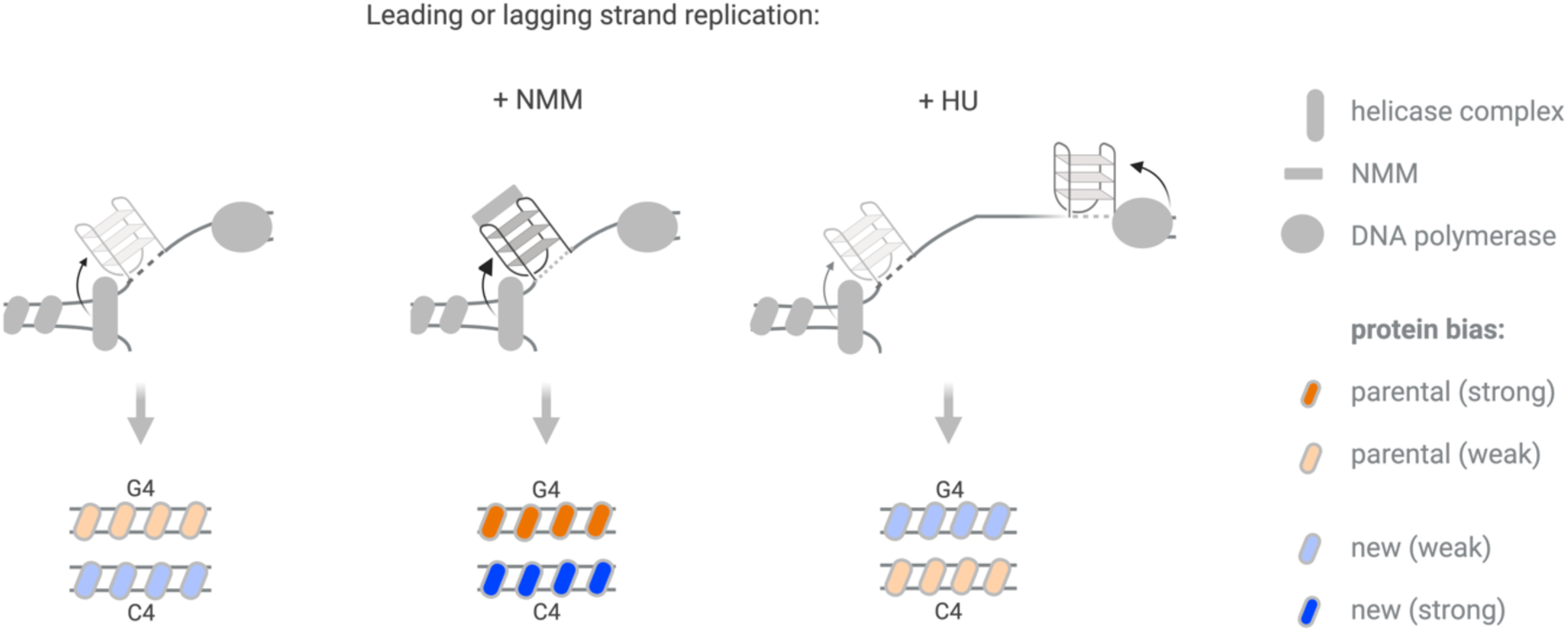
Asymmetric deposition of new and parental chromatin proteins at G4 sites during DNA replication. We suggest that the effect of G4s on the distribution of chromatin proteins may depend on how the G4 is detected by the cell. In both untreated cells (left panel) and cells treated with NMM (increasing the number of G4 sites, middle panel), G4 formation on either strand during DNA replication is immediately recognized and acted upon by the helicase complex. Among the consequently initiated processes likely is the recruitment of one or more specialized G4 helicases. We have shown that parental histones preferentially associate with the G4 strand in native and G4 stabilizing conditions, which suggests that there may also be a role for parental histone proteins in G4 resolution, or in maintaining the unresolved state of G4 sequences after resolution by a G4 helicase. In cells treated with HU (right panel), the bias in new chromatin protein deposition inverts towards the G4 strand. We propose that as a consequence of helicase-polymerase decoupling, ssDNA in HU-treated cells can form G4s that are not detected by the replicative helicase complex because it has already moved along. These G4s may only be recognized once the polymerase bumps into it, triggering an alternative response cascade with a different bias as a result.

Among the consequently initiated processes likely is the recruitment of one or more specialized G4 helicases such as DDX11 or FANCJ (18, 24). Our results show that parental histones preferentially associate with the G4 strand in native and G4-stabilizing conditions, suggesting that there may also be a role for parental histone proteins in G4 resolution, or in maintaining the unresolved state of G4 sequences after resolution by a G4 helicase. Finally, one cannot rule out the possibility that histones with parental PTMs simply have a greater affinity to DNA conformation in the vicinity of a formed quadruplex.

A previous study has shown that G4 presence can potentiate epigenetic instability in cells treated with a low dose of HU (7). HU is known to inhibit DNA synthesis by depleting nucleotides through the inhibition of ribonucleoside diphosphate reductase (25, 26). It has been suggested that the consequently slowed polymerases can get uncoupled from the helicase that unwinds the double-stranded DNA, leading to the increased exposure of single-stranded DNA ahead of the polymerases (7). Although these stretches of single-stranded DNA may get quickly coated with RPA, it is still conceivable that G4-formation is enhanced in these cells. We hence expected that treatment with HU, like treatment with NMM, would increase the deposition bias at G4 sequences. However, we find that HU treatment alone skews the distribution of new chromatin protein towards DNA that can form a G4. Thus, instead of strengthening the new protein deposition bias observed earlier, HU treatment inverts it. We suggest that ssDNA strands in HU-treated cells can form G4s that are not detected by the CMG because the helicase is already too far ahead (Figure 7). Therefore, the deposition of new chromatin protein at these G4s is simply skewed towards the strand that is replicated last, i.e. the strand that contains the replication-blocking G4.

It is worth noting that the reversal is not apparent for parental chromatin proteins as evidenced by the SCAR-seq results. This can be explained by a lower magnitude of the signal in the SCAR-seq method, asymmetric distribution of new and parental proteins upon fork slow-down, differential signals from histones and other chromatin proteins, or a combination thereof.

Rather unexpectedly, deposition of parental and new chromatin proteins observed upon combined NMM+HU treatment—which stabilizes G4s and potentially decouples replicative helicase from the replication fork—do not mirror each other (Figure 3C, D). After this treatment, the new chromatin proteins strongly avoid template G4 structures (like seen in NMM treatment), while old chromatin proteins do not show a strong alteration of their preference (like seen in HU treatment). This might mean that under such circumstances, sites with DNA replication blocks will lack some newly synthesized proteins, potentially resulting in the depletion of local protein occupancy. Finally, we can think of two non-mutually exclusive reasons why combined treatment with NMM and HU does not result in bias inversion. The first reason is that G4 stabilization increases the chance that G4s will form as soon as the DNA becomes single-stranded. Thus, most G4s will form near the CMG. This means that most G4s are detectable by the helicase in these cells, in contrast to cells that were only treated with HU.

The second reason is that G4 stabilization might slow the helicase to such an extent that helicase-polymerase uncoupling no longer occurs. A study in yeast indeed found that NMM treatment slowed cell growth and that cells accumulated in S-phase (27). We imagine that as there are more stable G4s forming in NMM-treated cells, the CMG is very frequently stalled at G4s on the leading strand, delaying DNA replication overall. Such slowing of the helicase would allow the DNA polymerases to keep up more easily, despite the HU-induced depletion of nucleotides. Thus, G4 stabilization could limit the exposure of single-stranded DNA behind the CMG. This would again mean that most G4s form near the CMG, where they are likely detected. The local effect of G4s on redistribution is evident from our fine-mapping analysis (Figures 4, 5) where we observe the strongest bias encompassing about two hundred base pairs around G4s, slightly shifted upstream. Our double-click-seq and SCAR-seq DNA fragment density analysis shows a strong phasing at G4 sites. The pattern is similar to that of nucleosome phasing, which hints that nucleosomes might be the direct effectors of biased protein redistribution at G4 sites.

Finally, global chromatin protein distribution patterns might, to some degree, be shaped by the distribution of G4 structures between the leading and lagging DNA strands. In general, as evidenced by our G4-Seq results, the quadruplexes are more often found on the lagging strand, where they are potentially less obstructive for replication. However, for a subset of G4 motifs with shorter loops, we see an increasing trend for leading strand preference. Assuming that G4s with small loops are more easily formed, our genome-wide survey suggests that they are more frequently found on the leading strand, while longer-loop motifs that are potentially corresponding to less prominent and unstable G4s are more often positioned to interfere with lagging strand replication. Accounting for these effects, as well as for other genomic features (e.g. different non-B DNA elements), will further improve our understanding of chromatin protein redeposition.

While there are still many unresolved questions about the role that G4 play in chromatin redistribution, it is clear that they underlie local disturbances in the symmetric inheritance of chromatin proteins. DNA quadruplexes can thus serve as prominent hot spots for epi-mutations and, genome-wide, be a major source behind stochastic changes in expression programs.

## Limitations

To determine the predominant replicating strand at the G4 sequences we used replication profiles of cells generated with OK-seq. We have previously compared the OK-seq replication profiles of cells grown with and without HU as we considered that HU treatment might affect the replication initiation zones, but we found them to be similar (8). Based on that we assume that the replication profiles are stable upon the treatments we used in this study.

## Materials and Methods

### Double-click-seq

We used hRPE-1 cells for our experiments. Double-click-seq experiments with cells grown in 1 μM NMM and a combination of 1 μM NMM and 200 μM hydroxyurea were done in triplicate and duplicate, respectively. In addition to our previously generated libraries, we generated one extra replicate for cells grown with hydroxyurea and without hydroxyurea. Experiments were done as previously described in (8). Briefly, *de novo* synthesized proteins and nascent DNA were labeled with respectively the methionine surrogate azidohomoalanine (AHA) and 5-Ethynyl-2′-deoxyuridine (EdU) for 24 hours after which nuclei were isolated. AHA-labeled histones in complex with DNA and other AHA-labeled proteins were enriched and subsequently, the protein-bound DNA was extracted and ligated to forked adaptors. Finally, the EdU-labeled nascent DNA was captured and the unlabeled parental DNA was eluted and amplified to generate a double-click-seq library. The resulting libraries were amplified using the NEBNext® Ultra™ DNA library prep kit for Illumina® using NEBNext® multiplex oligos for Illumina® and sequenced on a NextSeq 500 with onboard clustering using a 75-cycle high output v2.5 kit (Illumina) to generate paired-end 41-bp-long reads, as described previously in (8).

### SCAR-seq

We used hRPE-1 cells for our experiments. SCAR-seq experiments with cells grown in standard medium (untreated), 200 μM hydroxyurea, 1 μM NMM, and a combination of 1 μM NMM and 150 μM hydroxyurea were done in quadruplicate. After harvesting cells, SCAR-seq libraries were prepared following the previous published protocol (11) without any modification except using NEBNext® Ultra™ II DNA library prep kit (NEB) for the end repair, A-tailing, and adapter ligation of DNA samples. The library amplification was done with the same kit using NEBNext® multiplex oligos for Illumina® to label the samples. The quality of libraries was checked by an Agilent 4200 TapeStation System running either with Agilent D5000 or high sensitivity D5000 screen tape. The libraries with a bright band between 270bp to 300bp were sequenced on a NextSeq 500 with onboard clustering using a 75-cycle high output v2.5 kit (Illumina) to generate paired-end 41-bp-long reads, as described previously in (8).

### Raw data processing of double-click-seq and SCAR-seq data

Raw data processing of double-click-seq and SCAR-seq data was done as previously described for double-click-seq data (8). The quality of sequencing data and potential contaminations were evaluated by FastQC software (v. 0.11.5). Quality trimming was performed using paired-end mode implemented in the Trimmomatic package (v. 0.33). Trimmed sequences were mapped to genome assembly GRCh38 using Bowtie2 mapper (v. 2.2.4) with default parameters. Sorted alignments were marked for PCR duplicates using the Bamtools utility.

Data have been deposited to EBI ArrayExpress under accession numbers E-MTAB-8624 and E-MTAB-13346.

### Mass-spectrometry analysis of AHA labeling of proteins after the first (EdU) click reaction

The protocol was adapted from the double-click-seq protocol. Briefly, four T175 flasks were seeded with 3x10^6^ cells per flask with the complete culture medium. The next day, the culture medium was replaced by 95% AHA medium containing 0.2 mM AHA and 20 μM EdU. After 24 hours of incubation, cells were harvested by trypsinization. The cell pellet was washed twice with ice-cold PBS and stored at -80°C until further use. For the nuclei isolation, the cell pellet was resuspended in 1 ml of lysis buffer (0.1% NP-40, 1x proteinase inhibitor cocktail in PBS). The cell suspension was vortexed for ten seconds at maximum speed. The nuclei were pelleted by centrifugation for 10 seconds at 9000g. The nuclei were washed once with lysis buffer to remove membrane debris. Mono-nucleosomes were generated by MNAse digestion. Next, the Edu-click reaction was performed. The reaction components were added as follows, 1 μl of 100 Mm biotin-PEG4-azide, 10 μl of pre-mixed CuSO_4_ (5 μl, 50mM) and THPTA (5 μl, 250mM), 10 μl of freshly prepared 100 mM sodium ascorbate. The sample was mixed well and placed in an end-over-end rotator at room temperature for 45 minutes. The control group added all click reaction components except biotin-PEG4-azide. The labeled nucleosomes were enriched by Dynabeads^TM^ Streptavidin T1 following the manufacturer’s instruction.

Protein levels were determined with discovery-based proteomics (using label-free quantification) for relative protein concentrations (28). Briefly, in-gel digestion was performed on 40 µL of the proteins that were eluted from the beads in 1X LDS loading buffer (Thermo scientific) using trypsin (150 ng sequencing grade modified trypsin V5111; Promega) after reduction with 10 mmol/L dithiothreitol and alkylation with 55 mmol/L iodoacetamide proteins (29).

Discovery mass spectrometric analyses were performed on a quadrupole orbitrap mass spectrometer equipped with a nano-electrospray ion source (Orbitrap Exploris 480, Thermo Scientific) injecting 40% of the sample digests per measurement. Chromatographic separation of the peptides was performed by liquid chromatography (LC) on an Evosep system (Evosep One, Evosep) using a nano-LC column (EV1137 Performance column 15 cm x 150 µm, 1.5 µm, Evosep; buffer A: 0.1% v/v formic acid, dissolved in milliQ-H_2_O, buffer B: 0.1% v/v formic acid, dissolved in acetonitrile). Peptides were separated using the 30SPD workflow (Evosep). The mass spectrometer was operated in positive ion data-dependent acquisition mode (DDA) with the FAIMS unit switching between CV-45V and -60V using a cycle time of 1 sec per CV. MS spectra were acquired at a resolution of 120.000 at m/z 200 over a scan range of 375 to 1200 m/z with a normalized AGC target at 300% and an automatic maximum injection time mode. Peptide fragmentation was performed with higher energy collision dissociation (HCD) using a normalized collision energy (NCE) of 30%. The intensity threshold for ions selection was set at 8.0e3 with a charge exclusion of 1≤ and ≥6. The MS/MS spectra were acquired at a resolution of 15.000 at m/z 200, a normalized AGC target of 200%, a maximum injection time of 75 ms, and the isolation window set to 2 m/z.

### Proteomics data processing

Raw Thermo files (.raw) were converted using msconvert from the Proteowizzard toolset (downloaded on December 21, 2021) to mzML format in centroid format. The converted files were submitted to spectrum, peptide, and protein identification using MSFragger (version 3.4) and Philosopher (version 4.1.1). For MSFragger analysis the default settings for the closed search were used with the exception to include a variable modification for cysteine with a mass shift of -4.98632 Da. Swissprot downloaded on December 10, 2022 were used for closed search with MSFragger. After the search, the raw.pepxml files were further filtered using peptideProphet and proteinProphet of Philosopher for 1% FDR at PSM, peptide and protein levels. The search results were exported in tsv text format for PSM, peptide and protein level identifications.

### Raw data processing of OK-seq data

Quality trimming of raw OK-seq reads for RPE-1 hTERT cells (GSM3130725) was performed using single-end mode implemented in the Trimmomatic package (v. 0.36) using recommended settings and a minimum read length after trimming - 20bp. Trimmed sequences were mapped to genome assembly GRCh38 using Bowtie2 mapper (v. 2.2.4). Sorted alignments were marked for PCR duplicates using the Bamtools utility.

For the calculation of the effect of G4 loop size on the proportion of G4s on leading and lagging strands, we used scoring of Watson and Crick reads supplied with original data submission (GSM3130725 downloaded from GEO database and based on mapping to GRCh37 human genome reference). G4 regions were extracted from G4Seq submissions (GSM3003539) or predicted from GRCh37 sequence using nucleotide pattern (G_3+_N_1-x_G_3+_N_1-x_G_3+_N_1-x_G_3+_) with n ranging from 1 to 12 (tables deposited). G4 was considered predominantly located on the leading (or lagging) strand if the closest OK-seq datapoint was within 1 kb and had a Watson (Crick) score exceeding 0.2.

### Calculation of parental chromatin protein distribution bias at replication initiation zones

We calculated the average SCAR-seq signal across 9,608 previously identified autosomal replication initiation zones as described before for double-click-seq data (8) and plotted it. The signal was computed per 1 kb bins around each replication initiation zone, as the percentage of unambiguously mapped read pairs that correspond to (nascent) forward strand fragments. Fragments were assigned to bins according to the mapping coordinate of their central points. Read pairs with mapping quality below 20, discordantly mapped read pairs, and read pairs corresponding to fragments shorter than 145 bp apart (that might not originate from a nucleosome-bound fragment) were excluded. The signal was averaged across initiation zones. When combining results from replicates for Figure 1A, averaging was first applied across initiation zones and then across replicates.

We also visualized the SCAR-seq signal for selected individual replicates from untreated cells (replicate 2) and in cells treated with hydroxyurea (replicate 1) around each separate replication initiation zone in heatmaps with the R package pheatmap 1.0.12 (see Figure 1B and C).

### Calculation of the predominant replicating strand at G4 sequences on the template strand

Locations of experimentally defined G4 forming DNA sequences in the human genome in K^+^ conditions were downloaded from GEO accession GSE110582 in BED format. Coordinates were lifted from hg19 to hg38 assembly using the UCSC lift-over Genome Annotations tool with default settings. Only G4s on autosomes were used in our analyses (n= 412,112 out of a total of 433,618).

We determined the predominant replicating strand at these G4 sequences including flanks (2.5 kb to each side calculated from the G4 sequence center) using public OK-seq data from hRPE-1 cells. The OK-seq data contains single-end reads from Okazaki fragments. We, therefore, calculated the predominant replicating strand as the fraction of OK-seq reads that mapped to the G4 strand using the formula 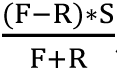, where F is the umber of reads mapped to the forward strand, R is the number of reads mapped to the reverse strand and S exhibits G4 strand, taking 1 for G4s on the forward strand and -1 for G4s on the reverse strand. A positive value indicates that replication at the G4 sequence on the template strand is mostly completed by leading strand replication, whereas a negative outcome indicates predominant lagging strand replication. We assumed that each read originated from an Okazaki fragment with a length of about 150 bp and used the mapping coordinate of the center of each fragment. We only considered primary alignments with a minimum mapping quality of 20, PCR duplicates were excluded. Scores for predominant replicating strands based on less than 20 reads were set to NA. We ended up with predominant replicating strand information for 390,129 potential autosomal G4 sequences (leading strand n=187,820, lagging strand n=196,796, and no preference n=5,513).

### Calculation of new and parental chromatin protein distribution at G4 sequences on the template strand

The new and parental chromatin protein distribution at potential G4 forming sequences was determined per 1 kb bins around G4 sequences that were grouped based on their potential to form parallel G4 structures (see the section below for details) or the predominant replicating strand at the sequence. The center of each G4 sequence corresponds to the center of bin “zero”.

Our double-click-seq data contains read pairs originating from the template strand of replicated DNA fragments bound to new chromatin proteins, where the strand of read 1 corresponds to the strand of the fragment. The bias in new chromatin protein deposition was therefore calculated as the percentage of unambiguously mapped read pairs of which read 1 mapped to the G4 strand. SCAR-seq data contains read pairs originating from the nascent strand of replicated DNA fragments bound to H4 histones with dimethylation at lysine 20 (H4k20me2, representing the parental chromatin proteins). Again, the strand of read 1 corresponds to the strand of the fragment, thus the bias in parental chromatin protein deposition was calculated as the percentage of unambiguously mapped read pairs of which read 2 mapped to the G4 template strand.

Read pairs were assigned to bins according to the mapping coordinate of the central points of the corresponding DNA fragments. Read pairs with mapping quality below 20, discordantly mapped read pairs, and read pairs corresponding to fragments shorter than 145 bp apart (that might not originate from a nucleosome-bound fragment) were excluded. To build the graphs in Figures 2, 3 and Supplementary Figures S4, and S5, the bias was averaged across G4 sequences. When combining results from replicates, averaging was first applied across G4s and then across replicates.

Further, we built heatmaps visualizing the bias at G4 sequences for selected, representative individual replicates. For the heatmaps, G4 sequences were sorted based on the predominant replicating strand at the G4 sequence (see section above for its calculation). To limit the number of rows to be visualized in the heatmaps, we averaged the redistribution bias per 100 G4 sequences. Bias averages were normalized per row by calculating their z-score, with red and blue indicating respectively increased and decreased chromatin protein deposition onto replicating strands that may be obstructed by a G4 on the template strand. Heatmaps were plotted with the pheatmap R package.

We also calculated the new and parental protein distribution per position within 1kb to each side of potential G4 forming sequences. The center of each G4 sequence corresponds to position “0”. Read pairs were assigned to positions according to the mapping coordinate of the central points of the corresponding DNA fragments. Again, read pairs with mapping quality below 20, discordantly mapped read pairs, and read pairs corresponding to fragments shorter than 145 bp apart were excluded. For each double-click-seq replicate, the bias in new chromatin protein deposition was calculated as the percentage of unambiguously mapped read pairs of which read 1 mapped to the G4 strand. For each SCAR-seq replicate, the bias in parental chromatin protein deposition was calculated as the percentage of unambiguously mapped read pairs of which read 2 mapped to the G4 template strand. Next, percentages were averaged per 20 bp window after which they were averaged across replicates.

### Determining confidence intervals

For each double-click-seq and SCAR-seq replicate, 1000 bootstrap replicates were performed to assess the robustness of the observed effect at G4 sequences and replication initiation zones. We used the range between the 2.5th and the 97.5th percentiles to show 95% confidence intervals on the plots.

### Selection of potential parallel-stranded G4 sequences

G4 motifs with triple guanine stretches, two non-guanine one-nucleotide loops, and one loop between one and three bases (of which the middle can be guanine) can reportedly only form parallel G4s, whereas motifs with longer loops can also form hybrid and antiparallel G4s (22). We, therefore, selected autosomal G4 sequences that contained such a motif as parallel-stranded G4 sequences (n=15,010, with predominant replicating strand information available for n=14,348 of those).

### Calculation of new and parental chromatin protein occupancy profiles at G4 sequences and Transcription Start Sites

New and parental chromatin protein occupancy profiles at potential G4 forming sequences were determined per single base pair position within 1kb to each side of the G4 sequence / Transcription Start Site (TSS). TSSs (n=19912) were defined as the gene starts of all protein-coding genes reported in the human genome build GRCh38 annotation v.92 from Ensembl. The center of each G4 sequence / TSS corresponds to position “0”. Read pairs were assigned to positions according to the mapping coordinate of the central points of the corresponding DNA fragments. Again, read pairs with mapping quality below 20, discordantly mapped read pairs, and read pairs corresponding to fragments shorter than 145 bp apart were excluded. Counts were summed across replicates, converted to percentages (% fragments mapping to that position out of the total number of fragments mapping to the entire 2 kb region), and averaged per 20 bp window.

## Data Availability

Data have been deposited to EBI ArrayExpress under accession numbers E-MTAB-8624 and E-MTAB-13346. The code generated during this study is available at GitHub; https://github.com/thamarlobo/G4_chromatin_analysis.

## Supporting information

### Supplementary Figures

**Figure S1.**
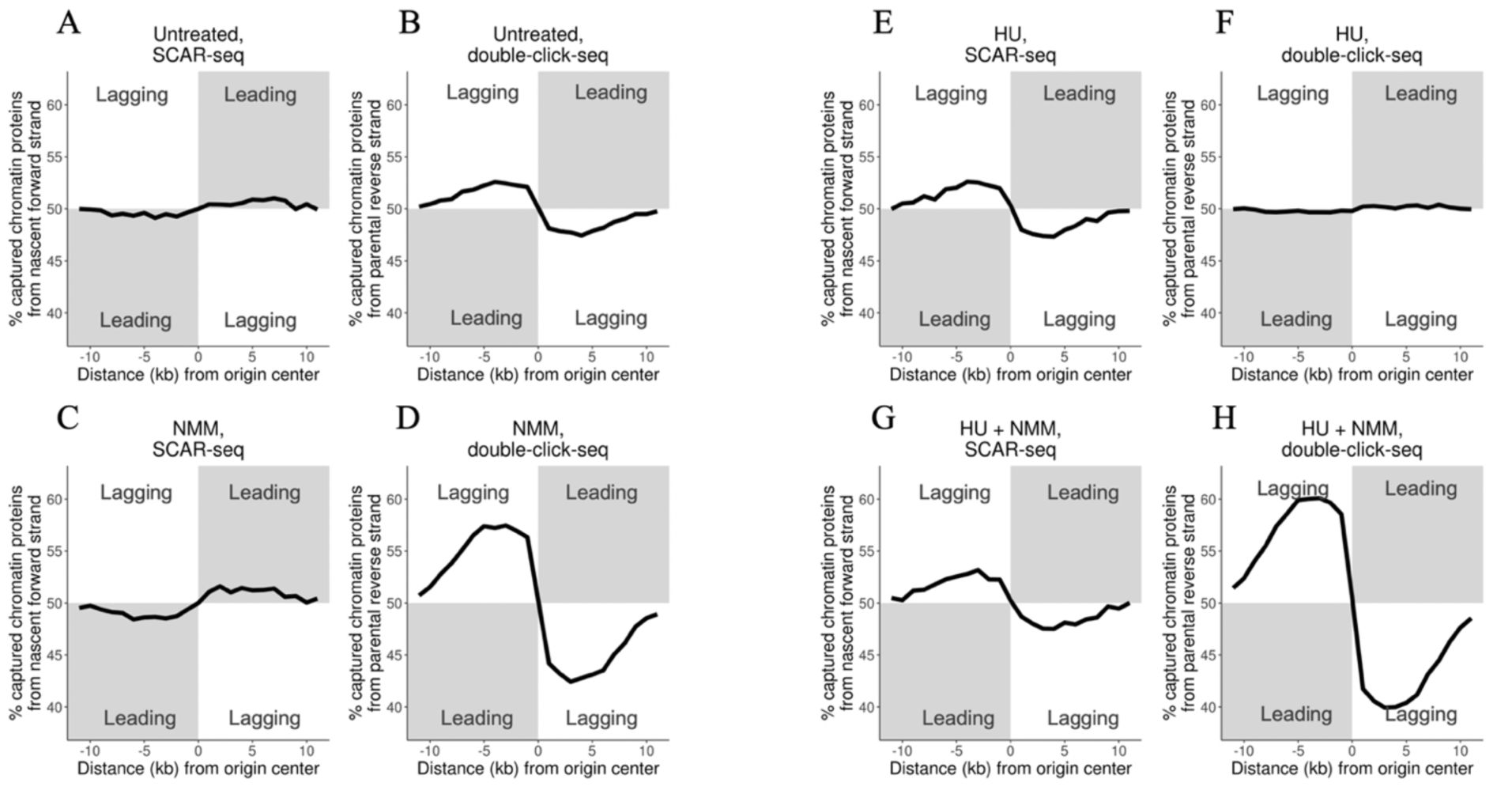
The distributions of parental and newly synthesized proteins at replication initiation zones mirror each other across experimental conditions, except for combined treatment (HU + NMM). A & B. Average bias of parental (left) and new (right) chromatin protein deposition at replication initiation zones in untreated cells. C & D. Average bias of parental (left) and new (right) chromatin protein deposition at replication initiation zones in cells treated with NMM. E & F. Average bias of parental (left) and new (right) chromatin protein deposition at replication initiation zones in cells treated with HU. G & H. Average bias of parental (left) and new (right) chromatin protein deposition at replication initiation zones in cells treated with both HU and NMM.

**Figure S2.**
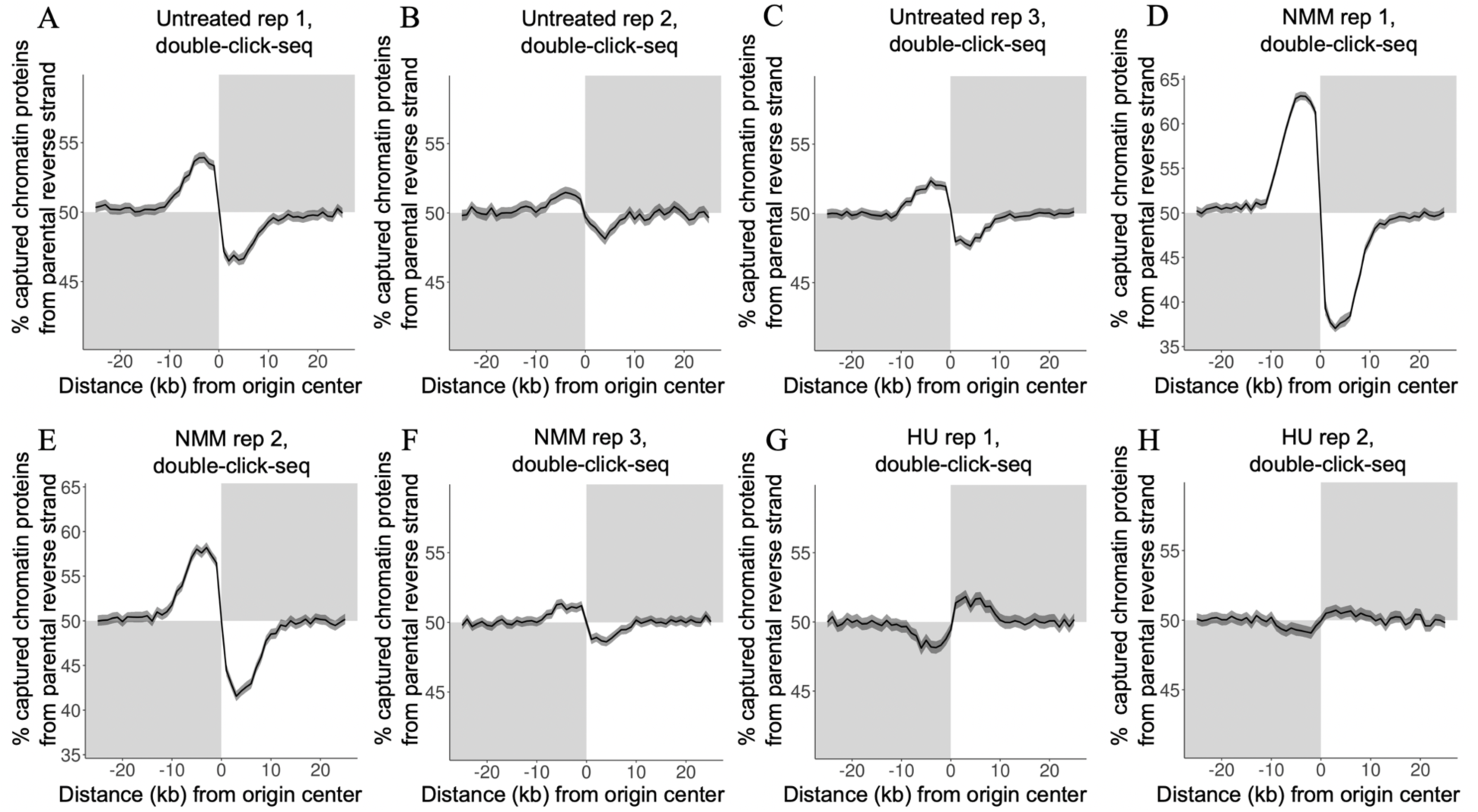
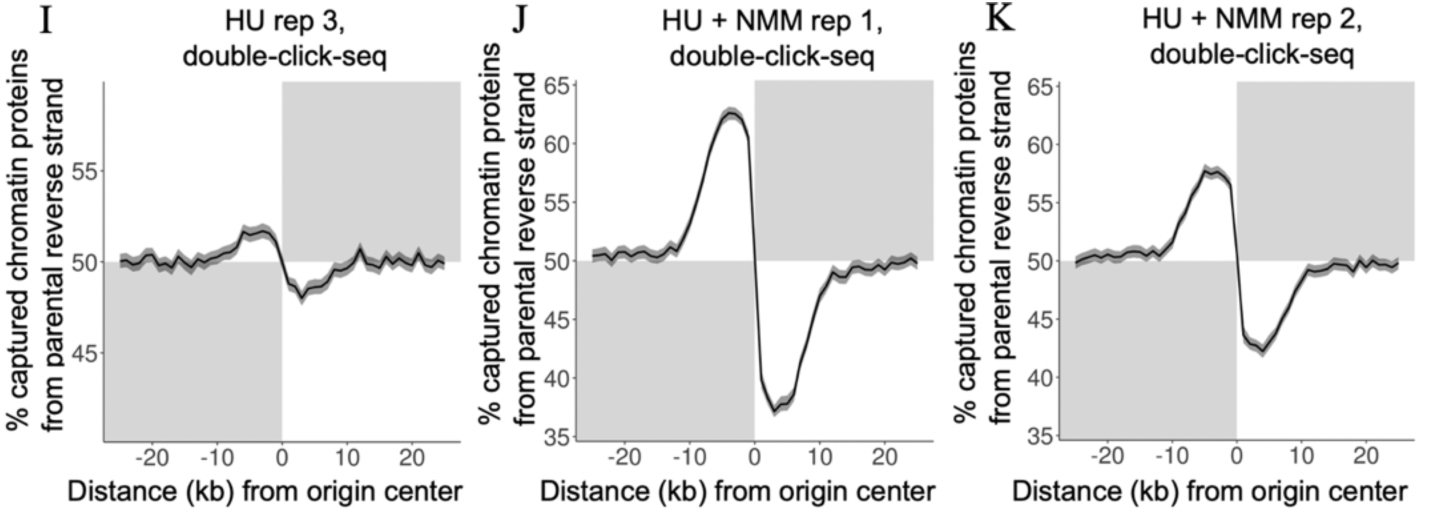
Average new chromatin protein deposition bias at replication initiation zones including 95% confidence intervals (light fill). The deposition was calculated as the percentage of double-click-seq reads that mapped to the reverse strand, out of the total number of double-click-seq reads mapping at that location. The treatment is indicated at the top of each panel.

**Figure S3.**
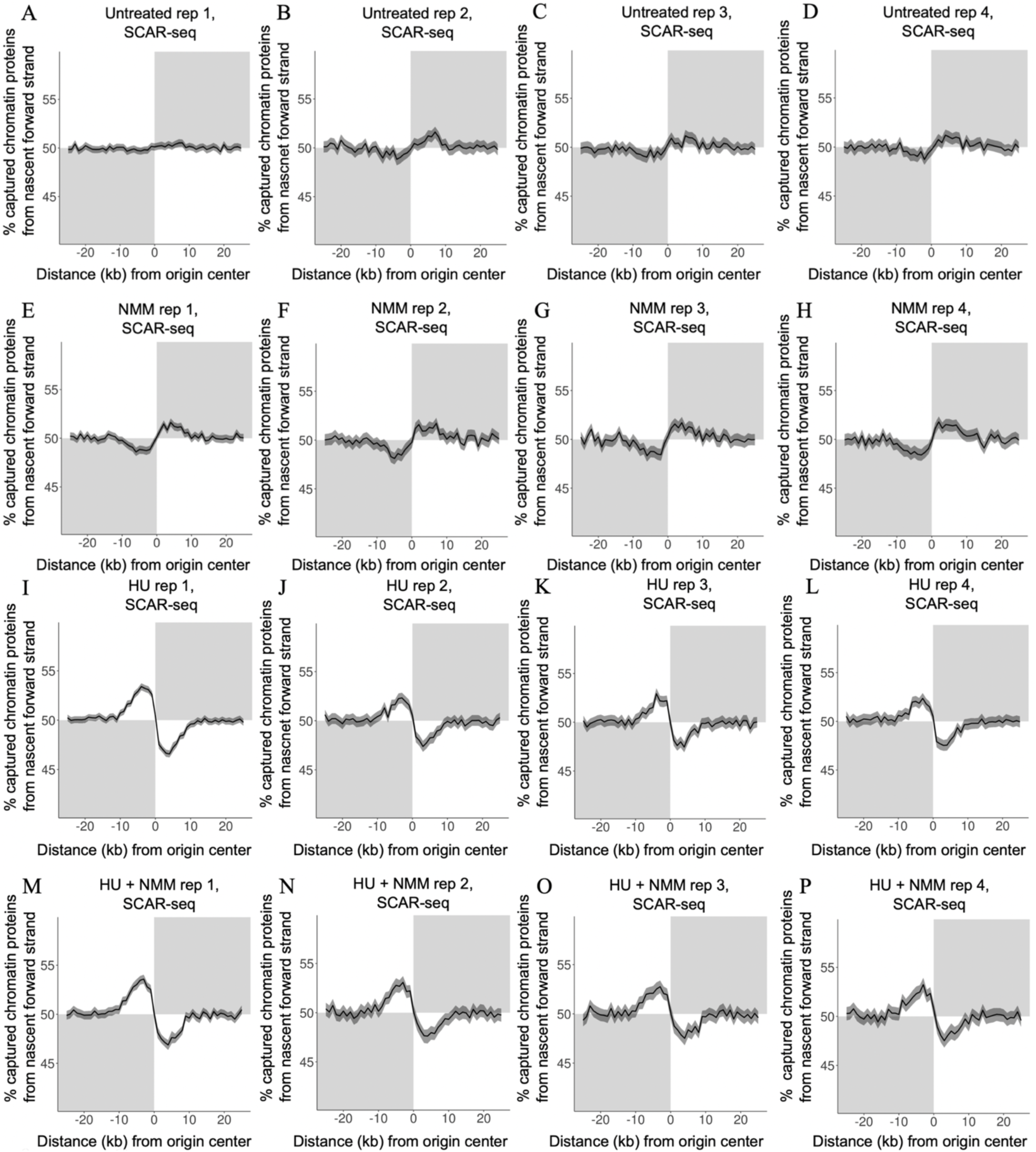
Average parental chromatin protein deposition bias at replication initiation zones including 95% confidence intervals (light fill). The deposition was calculated as the percentage of SCAR-seq reads that mapped to the forward strand, out of the total number of SCAR-seq reads mapping at that location. The treatment is indicated at the top of each panel.

**Figure S4.**
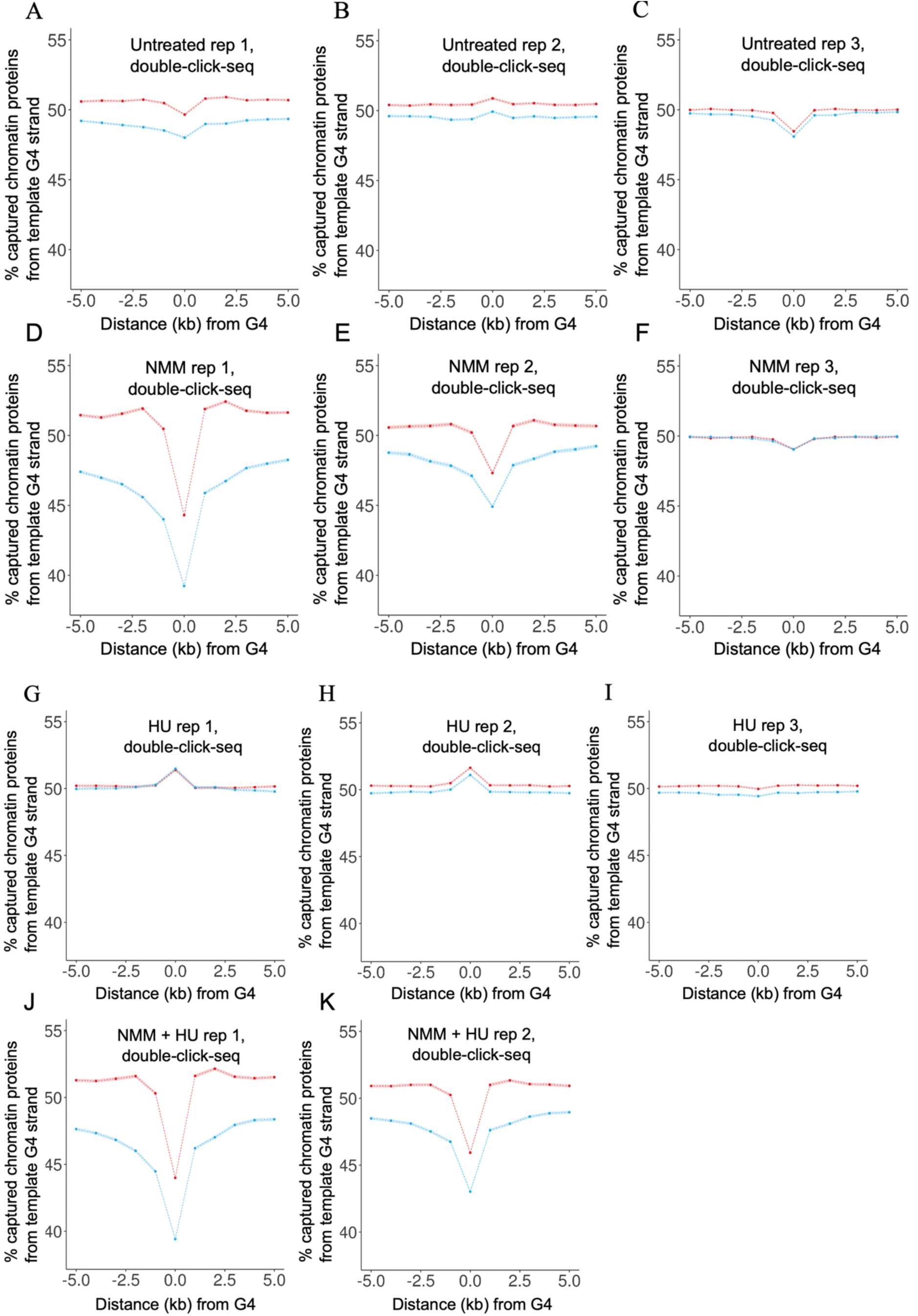
Average new chromatin protein deposition bias at G4 sequences including 95% confidence intervals (light fill). We determined the average deposition separately for G4 sequences that could interfere with predominant leading (red) and lagging strand replication (blue). The deposition was calculated as the percentage of captured chromatin proteins bound to the G4 template strand. The treatment is indicated at the top of each panel.

**Figure S5.**
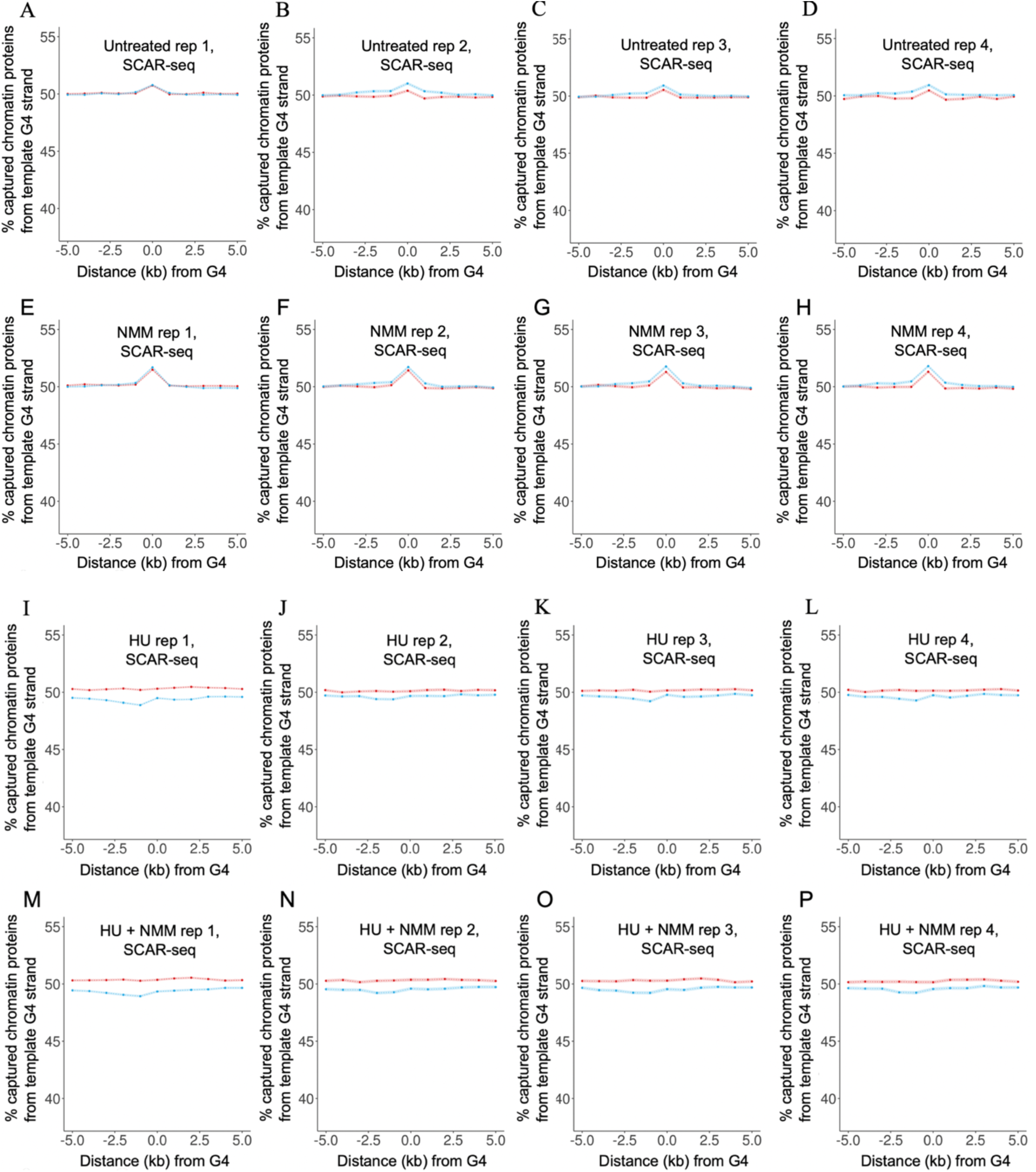
Average parental chromatin protein deposition bias at G4 sequences including 95% confidence intervals (light fill). We determined the average deposition separately for G4 sequences that could interfere with predominant leading (red) and lagging strand replication (blue). The deposition was calculated as the percentage of captured chromatin proteins bound to the G4 template strand. The treatment is indicated at the top of each panel.

**Figure S6.**
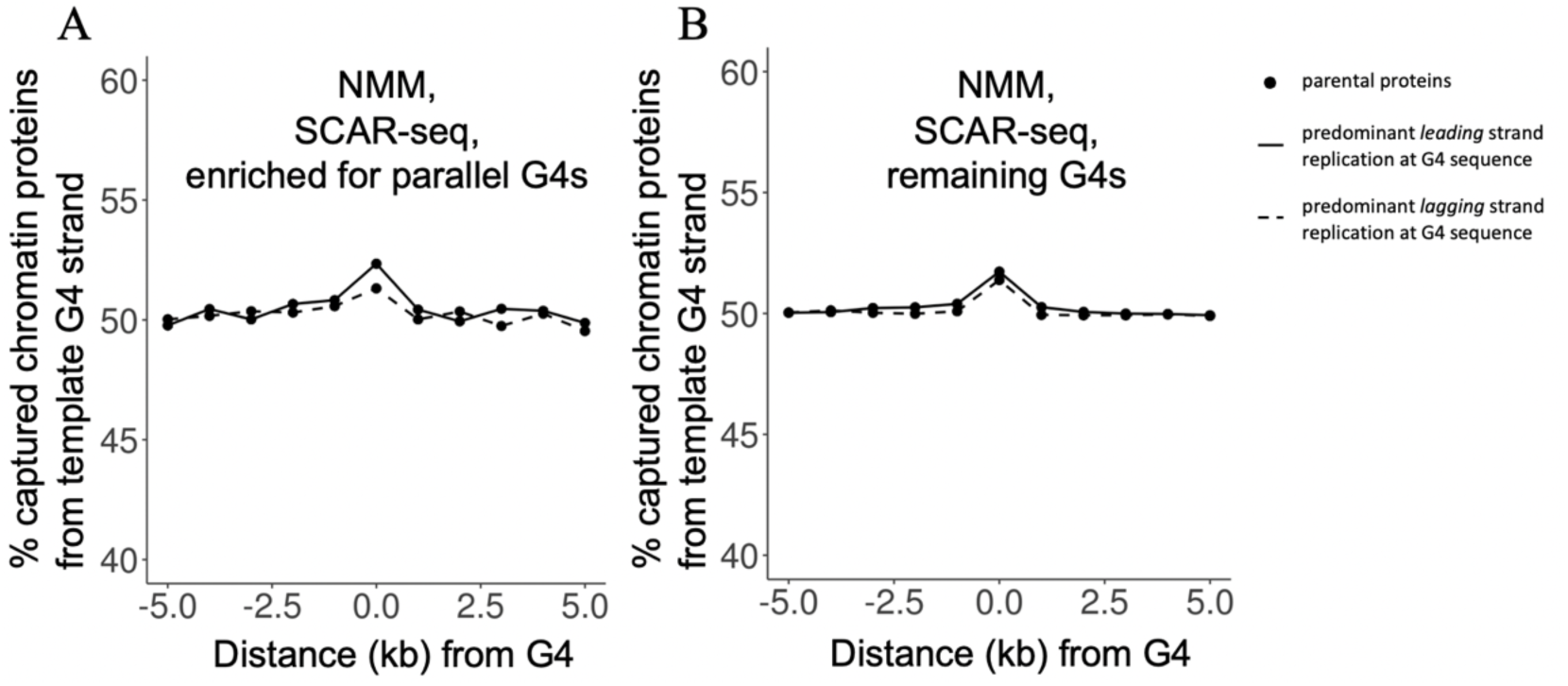
Average parental chromatin protein deposition at parallel and remaining G4 sequences. We determined the average deposition separately for G4 sequences that could interfere with predominant leading (continuous line) and lagging strand replication (dashed line). The deposition was calculated as the percentage of captured chromatin proteins bound to the G4 template strand. A.The average deposition of parental chromatin proteins at parallel G4 sequences in cells treated with NMM. B. The average deposition of parental chromatin proteins at remaining G4 sequences in cells treated with NMM.

**Figure S7.**
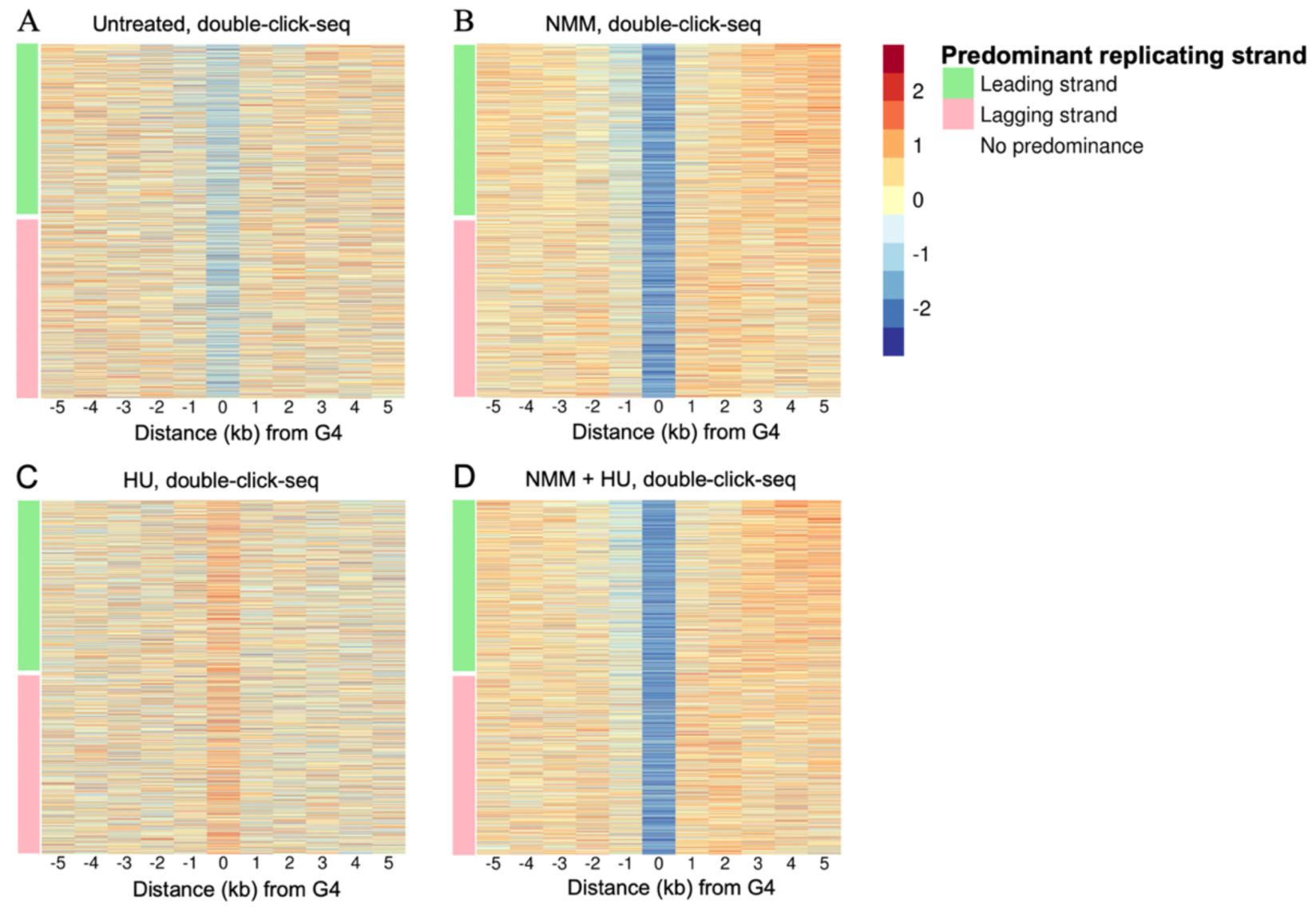
Heatmaps of bias of new chromatin protein deposition at G4 sequences. G4 sequences were sorted by predominant replicating strand at the G4 sequence (as calculated using OK-seq replication profiles). The percentage of captured chromatin proteins bound to the G4 template strand was determined per 1 kb bin around each G4 sequence. To limit the number of mapped rows, we averaged the bias per 100 G4 sequences. Bias averages were z-score normalized per row, with red and blue indicating respectively increased and decreased new chromatin protein deposition onto the template strand that can form a G4. For each row, the color bar on the left indicates the predominant replication type at the template strand that can form a G4: leading (green), lagging (pink), or no predominance (white). A. Bias in untreated cells, replicate 3. B. Bias in cells treated with NMM, replicate 1. C. Bias in cells treated with hydroxyurea, replicate 1. D. Bias in cells treated with NMM and hydroxyurea, replicate 1.

**Figure S8.**
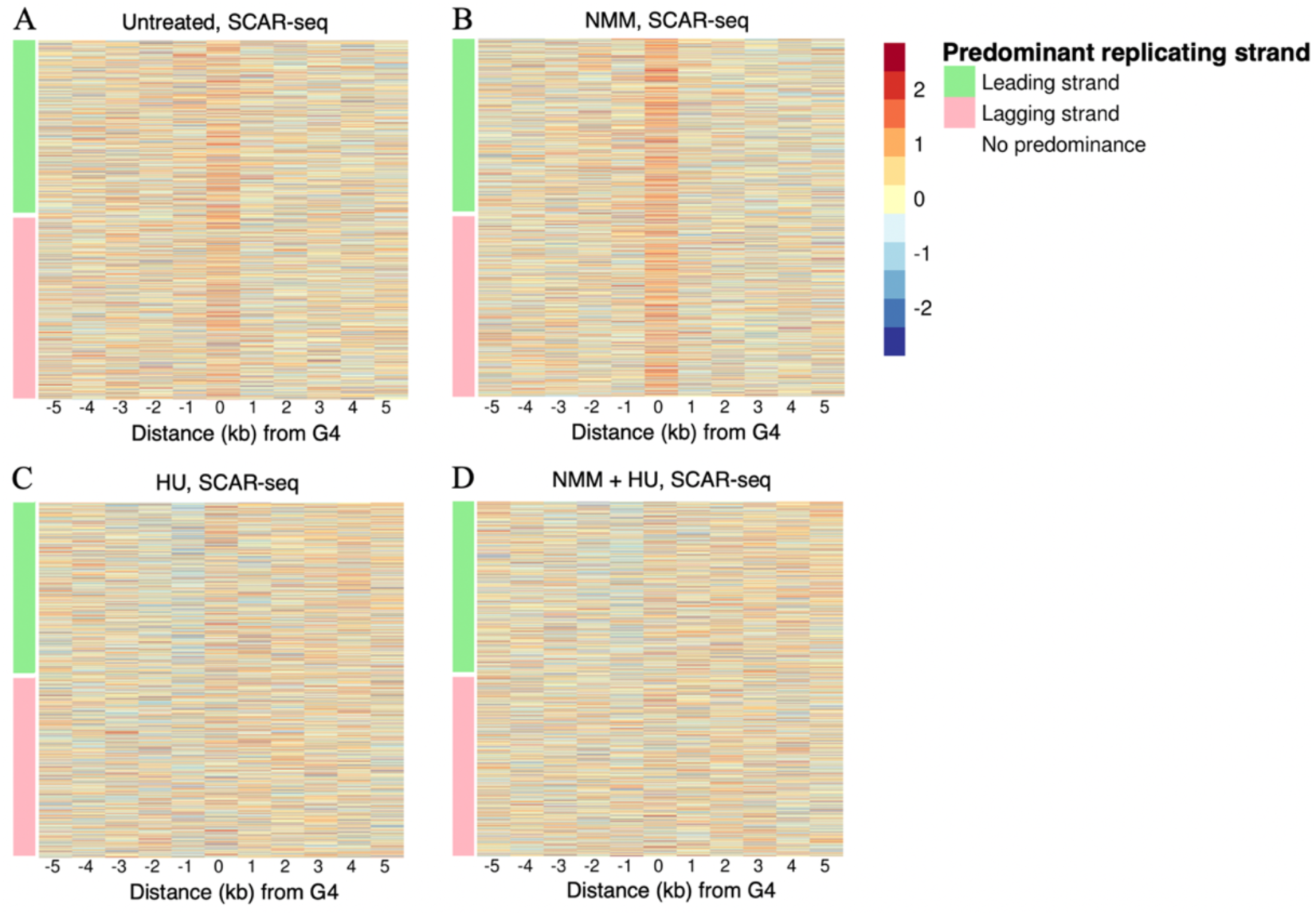
Heatmaps of bias of parental chromatin protein deposition at G4 sequences. G4 sequences were sorted by predominant replicating strand at the G4 sequence (as calculated using OK-seq replication profiles). The percentage of captured chromatin proteins bound to the G4 template strand was determined per 1 kb bin around each G4 sequence. To limit the number of mapped rows, we averaged the bias per 100 G4 sequences. Bias averages were z-score normalized per row, with red and blue indicating respectively increased and decreased parental chromatin protein deposition onto the template strand that can form a G4. For each row, the color bar on the left indicates the predominant replication type at the template strand that can form a G4: leading (green), lagging (pink), or no predominance (white). A. Bias in untreated cells, replicate 1. B. Bias in cells treated with NMM, replicate 1. C. Bias in cells treated with hydroxyurea, replicate 1. D. Bias in cells treated with NMM and hydroxyurea, replicate 1.

**Figure S9.**
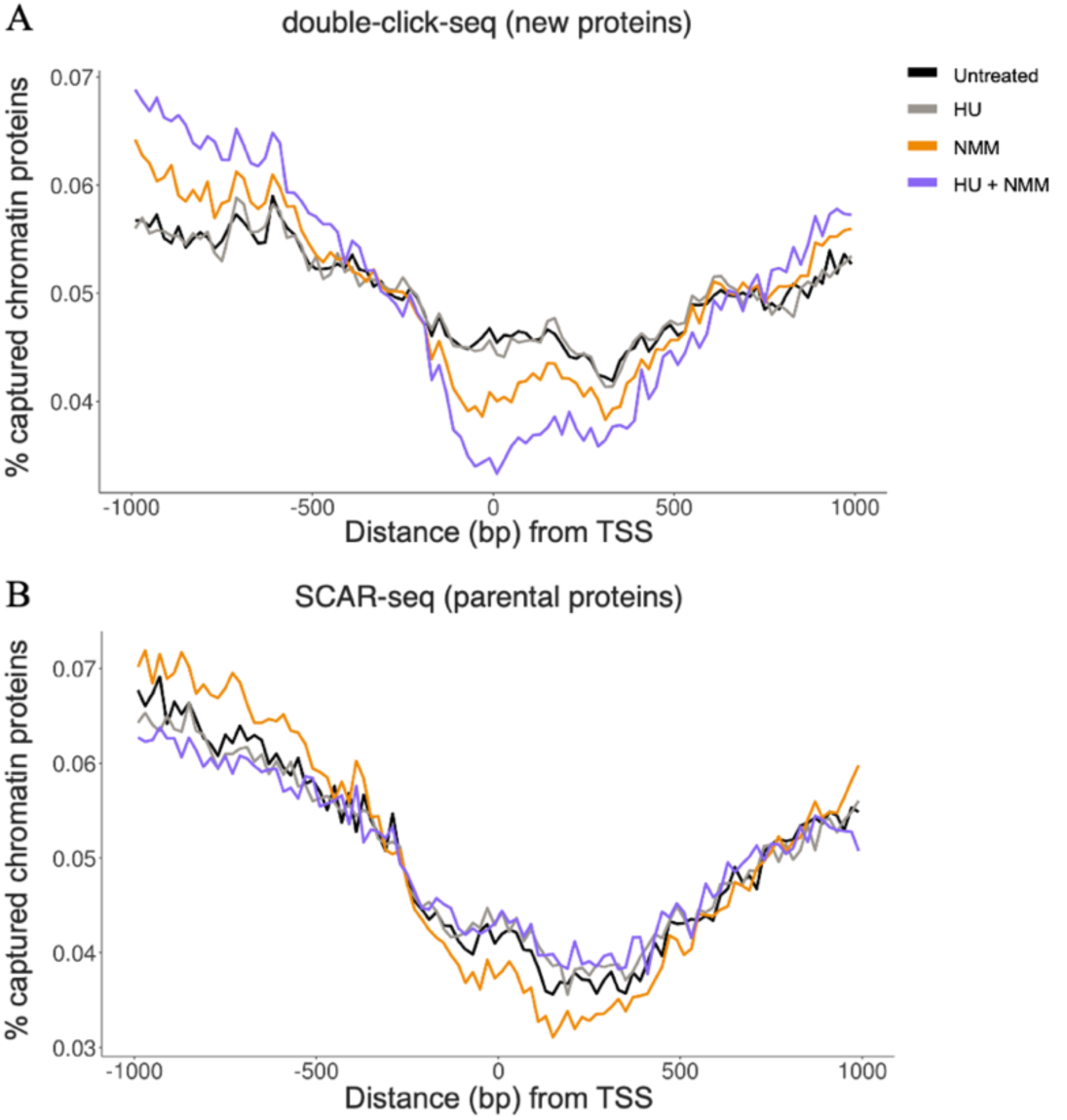
Average new and parental chromatin protein occupancy at Transcription Start Sites (TSS). The signal was summed across replicates, converted to percentages, and averaged per 20bp windows. A. The average occupancy of new proteins. B. The average occupancy of parental proteins.

**Table S1.** Peptide spectral matches (PSMs) significantly overrepresented in EdU-clicked (compared to no-click) mass-spec data. Two-tailed chi-square test was used to calculate difference in PSMs identified in EdU-click results (4,704 PSMs in total) as compared to no-click sample (4,054 total PSMs). Benjamini-Hochberg multiple testing correction procedure was used to calculate adjusted p-value (FDR)

| Protein | EdU-click, PSMs | No-click, PSMs | P-value | FDR |
| --- | --- | --- | --- | --- |
| H2AC17 | 130 | 12 | 7.25e-20 | 9.55e-17 |
| AHNAK | 130 | 17 | 1.57e-17 | 1.03e-14 |
| VIM | 115 | 14 | 4.01e-16 | 1.76e-13 |
| H2BC12L | 62 | 0 | 2.14e-14 | 7.03e-11 |
| PLEC | 59 | 0 | 8.13e-13 | 2.14e-10 |
| SPTAN1 | 53 | 0 | 1.18e-11 | 2.22e-09 |
| ACTB | 67 | 12 | 2.49e-08 | 4.11e-06 |
| ACTN4 | 57 | 10 | 2.30e-07 | 3.37e-05 |
| HNRNPUL1 | 27 | 0 | 1.34e-06 | 1.47e-04 |
| MYOF | 26 | 0 | 2.10e-06 | 1.98e-04 |
| SPTBN1 | 25 | 0 | 3.30e-06 | 2.72e-04 |
| FLNA | 31 | 3 | 1.12e-05 | 7.01e-04 |
| SLC3A2 | 21 | 0 | 2.03e-05 | 1.16e-03 |
| CKAP4 | 20 | 0 | 3.20e-05 | 1.69e-03 |
| ANXA6 | 20 | 0 | 3.20e-05 | 1.69e-03 |
| RPSA | 23 | 1 | 3.37e-05 | 1.69e-03 |
| HSP90B1 | 23 | 1 | 3.37e-05 | 1.69e-03 |
| ITGB1 | 24 | 2 | 7.63e-05 | 3.14e-03 |
| ACTA1 | 18 | 0 | 7.98e-05 | 3.19e-03 |
| RACK1 | 26 | 3 | 9.97e-05 | 3.75e-03 |
| SEPTIN11 | 16 | 0 | 2.00e-04 | 5.86e-03 |
| ATP1A1 | 16 | 0 | 2.00e-04 | 5.86e-03 |
| LMO7 | 15 | 0 | 3.18e-04 | 8.20e-03 |
| ALCAM | 15 | 0 | 3.18e-04 | 8.20e-03 |

**Table S2.**
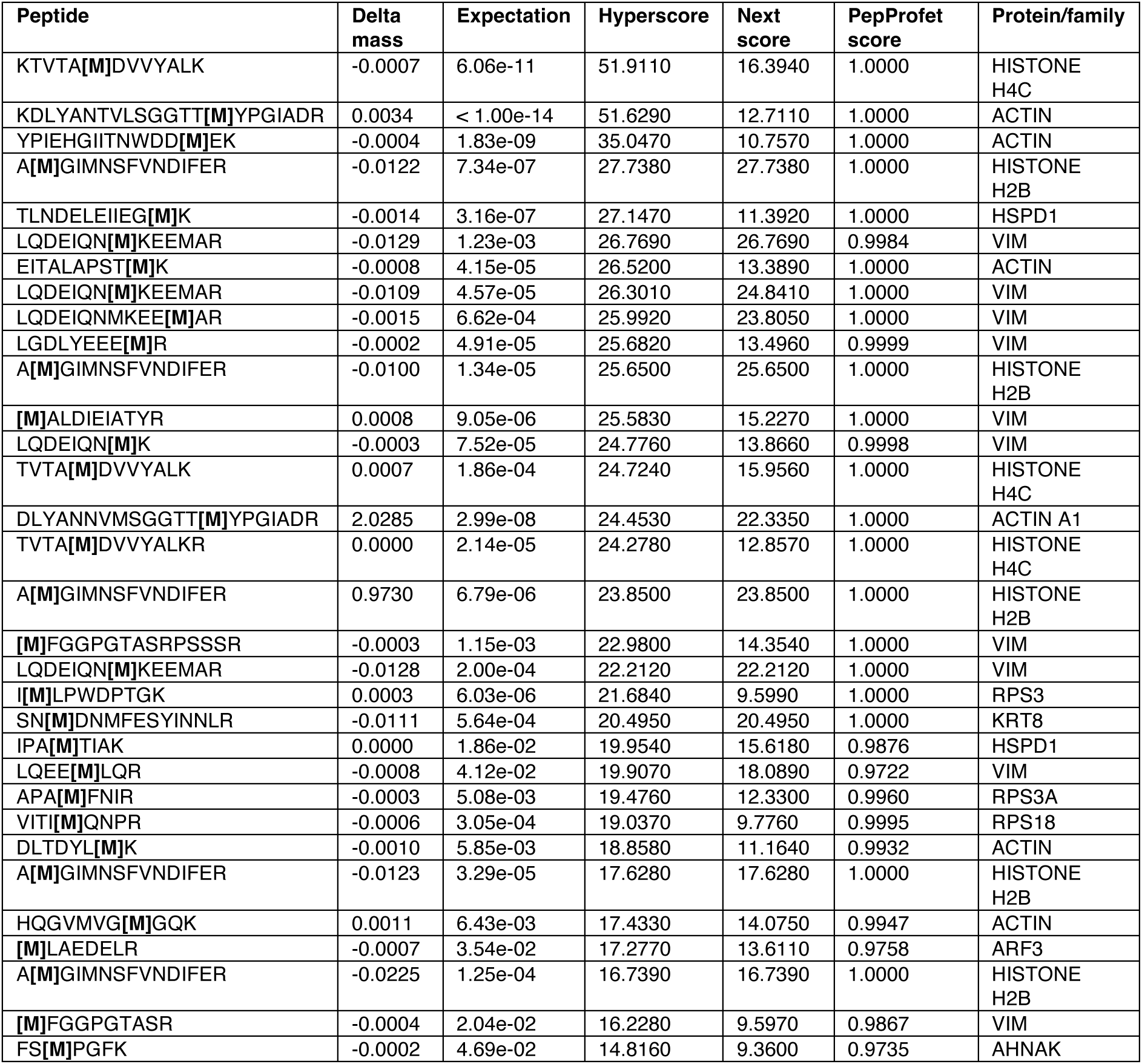
Modified peptides (Met->AHA, delta weight = -4.98632 a.u.) identified in EdU-click sample. The PSMs are ordered by hyperscore. Methionines that are predicted to be replaced by AHA are given in square brackets and bold font.

**Table S3.** Experimental condition, number of input, and mapped read pairs for each.

| Experimental condition | Library type | Number of input read pairs | Informative read pairs* |
| --- | --- | --- | --- |
| WT rep 1 | double-click-seq | 40,268,580 | 19,945,411 |
| WT rep 2 | double-click-seq | 40,733,126 | 15,373,997 |
| WT rep 3 | double-click-seq | 38,017,658 | 27,911,289 |
| 200 uM HU rep 1 | double-click-seq | 40,086,193 | 13,450,567 |
| 200 uM HU rep 2 | double-click-seq | 36,693,399 | 16,369,979 |
| 200 uM HU rep 3 | double-click-seq | 28,419,780 | 20,556,856 |
| 1 uM NMM rep 1 | double-click-seq | 39,742,799 | 20,406,022 |
| 1 uM NMM rep 2 | double-click-seq | 48,571,438 | 8,822,464 |
| 1 uM NMM rep 3 | double-click-seq | 34,726,058 | 25,458,954 |
| 200 uM HU + 1 uM NMM rep 1 | double-click-seq | 39,903,185 | 16,593,611 |
| 200 uM HU + 1 uM NMM rep 2 | double-click-seq | 38,422,981 | 12,331,423 |
| WT rep 1 | SCAR seq | 69,797,909 | 18,377,697 |
| WT rep 2 | SCAR seq | 34,041,832 | 8,288,757 |
| WT rep 3 | SCAR seq | 32,715,457 | 8,029,517 |
| WT rep 4 | SCAR seq | 33,283,073 | 8,118,347 |
| 200 uM HU rep 1 | SCAR seq | 89,088,658 | 23,347,569 |
| **200 uM HU rep 2** | SCAR seq | 34,007,316 | 9,426,928 |
| **200 uM HU rep 3** | SCAR seq | 33,497,919 | 9,385,243 |
| **200 uM HU rep 4** | SCAR seq | 31,775,172 | 8,937,882 |
| **1 uM NMM rep 1** | SCAR seq | 74,570,916 | 17,736,046 |
| **1 uM NMM rep 2** | SCAR seq | 33,736,613 | 8,649,802 |
| **1 uM NMM rep 3** | SCAR seq | 31,203,353 | 8,152,728 |
| **1 uM NMM rep 4** | SCAR seq | 32,845,685 | 8,471,769 |
| **150 uM HU + 1 uM NMM rep 1** | SCAR seq | 71,398,166 | 14,433,551 |
| **150 uM HU + 1 uM NMM rep 2** | SCAR seq | 31,639,213 | 6,796,506 |
| **150 uM HU + 1 uM NMM rep 3** | SCAR seq | 30,767,968 | 6,821,005 |
| **150 uM HU + 1 uM NMM rep 4** | SCAR seq | 28,612,700 | 6,257,508 |

\*concordantly and uniquely (mapq >=20) mapped to autosomes, PCR duplicates excluded, fragment length at least 145 bp.

